# Somatosensory Cortical Signature of Facial Nociception and Vibrotactile Touch Induced Analgesia

**DOI:** 10.1101/2022.04.14.488349

**Authors:** Jinghao Lu, Bin Chen, Manuel Levy, Peng Xu, Bao-Xia Han, Jun Takatoh, P. M. Thompson, Zhigang He, Vincent Prevosto, Fan Wang

**Affiliations:** Department of Brain and Cognitive Sciences, McGovern Institute for brain research, MIT; Department of Neurobiology, Duke University; Department of Psychology and Neuroscience, UNC; Kirby Neurobiology Center, Boston Children’s Hospital, Harvard Medical School

## Abstract

Pain relief by vibrotactile touch is a common human experience. Previous neurophysiological investigations in animals focused on spinal mechanisms while human studies suggested the involvement of supraspinal mechanisms. Here we asked whether and how the primary somatosensory cortex (S1) is involved in touch induced analgesia. We discovered that in mice, vibrotactile reafferent signals from self-generated whisking significantly reduce facial nociception, which is abolished by specifically blocking touch transmission from thalamus to the barrel cortex (S1B). The presence of whisking altered nociceptive signal processing in S1B neurons. Intrinsic manifold analysis of S1B population activity revealed that whisking pushes the transition of neural state induced by noxious stimuli towards the state encoding non-nocifensive actions. Thus, S1B integrates facial tactile and noxious signals to enable touch mediated analgesia.

**Teaser:** Vibrotactile signals modulate barrel cortex population dynamics during touch mediated facial analgesia

## Introduction

When a part of our body is hurt, we often instinctively rub, massage or shake it. This phenomenon is referred to as touch-mediated-analgesia, or pain relief by touch. It prompted the original development of the gate control theory (*1*). Subsequently, neurophysiological studies in anesthetized animals identified a class of wide dynamic range (WDR) neurons in the spinal cord, which receive both touch and pain inputs in the dorsal horn, as the likely substrate for the analgesia mediated by touch. The receptive field of WDR neurons has a center-surround concentric arrangement: the center is *activated* by tactile or nociceptive stimuli whereas the surround is activated by noxious stimuli and *inhibited* by touch stimuli. Thus, vibrotactile stimuli applied around the noxious stimulus-sensitive center reduce WDR neuronal responses to the nociceptive inputs through inhibition (*2–4*). In humans, touch mediated pain relief has been demonstrated for both the body parts innervated by dorsal root ganglia (DRG) neurons, as well as for orofacial regions innervated by trigeminal ganglion (TG) neurons (*5*). A quantitative study in humans, in which a painful laser stimulation was flanked by two touch stimuli at varying distances, showed that concurrent application of touch caused the subjects to rate the laser stimuli as less painful in a distance dependent manner, consistent with the receptive field properties of spinal WDR neurons (*6*). Interestingly, the same study also found that although the “pinprick laser pain” sensation was reduced by concurrent touch stimuli, the latencies to detect the laser stimuli were not affected (i.e. noxious signals detected by Aδ fiber were relayed to the brain). This observation suggested that supraspinal mechanisms are involved in the “interpretation” of the laser stimuli as “less/non-painful” (*6*). Other human studies suggested a possible cortical involvement in this process (*7–9*). For example, tactile stimulus delivered 60ms after the application of noxious electrical stimulus whose signal has already reached the cortex by 60ms, can still inhibit cortical responses to pain (*7*). However, if there is a cortical mechanism, which cortical areas would be involved remains unclear. Answering this question requires a reliable awake behaving animal model of touch-mediated analgesia.

It is well known that vibrotactile information is mainly processed by the primary somatosensory cortex (S1). Abnormal activation of S1 was linked to mechanical allodynia in animal pain models (*10, 11*). Numerous human imaging and electroencephalography EEG studies also revealed that S1 is activated by painful stimuli (*12–14*). A recent study further showed that nociceptive stimuli elicited strong gamma-band oscillations in the superficial layers of S1 (*15*). Thus, S1 has the potential to integrate tactile and nociceptive information. In rodents, it has long been known that the S1 barrel cortex (S1B) is dedicated to process whisker derived tactile information such as the location, size, and texture of objects touched by the whiskers (*16–22*). By contrast, little is understood about whether and how S1B processes nociceptive information derived from the whisker pad skin. To develop a rodent orofacial model of touch mediated analgesia, we hypothesized that whisking, as a form of vibrotactile inputs, might naturally suppress the nociception of acute noxious stimuli experienced by the face, and if so, this would provide a model to examine S1B’s role in touch, nociception, and their interactions. In this study, we tested our hypothesis and discovered that self-initiated whisking could indeed suppress the mice’s nocifensive responses, and this effect was diminished when the transmission of whisking tactile signal from ventroposterior medial nucleus (VPM) in the thalamus to S1B was blocked. We further characterized the nociceptive representation in the layer 2/3 (L2/3) neurons of S1B, and uncovered potential population activity patterns underlying whisking induced suppression of nociception.

## Results

### Reafferent signals from self-generated whisking suppress facial nociception

The first question we asked was whether vibrotactile afferent signals of self-generated whisking would suppress orofacial nocifensive responses to noxious heat and mechanical stimuli in mice. We habituated head-fixed mice (C57BL/6J, n = 9, with 5 male and 4 female) on the running wheel and applied either radiant heat or noxious mechanical von Frey filament stimuli to the left whisker pad of the mice. Head-fixation allowed consistent targeting of stimuli to the whisker pad (Movie S1). Depending on the intensity of noxious stimuli they perceived, mice would either wipe their face, a nocifensive behavior, or not do so (Fig. 1A, Movie S1). We operationally used this face wiping behavior as a surrogate for nociceptive perception. We applied the stimuli when mice were either whisking-in-air (and running), or resting (no-whisking) (Fig. 1A). Interestingly, we found that the number of face-wiping in response to noxious heat or mechanical stimuli was significantly reduced when mice were whisking, compared to when mice were not whisking (Fig. 1B, p < 0.025 * or p < 0.01 **, when applicable). Similarly, the percentage of the trials in which mice wiped their face was also significantly reduced under conditions of self-generated whisking (Fig. S1A).

**Fig.1.**
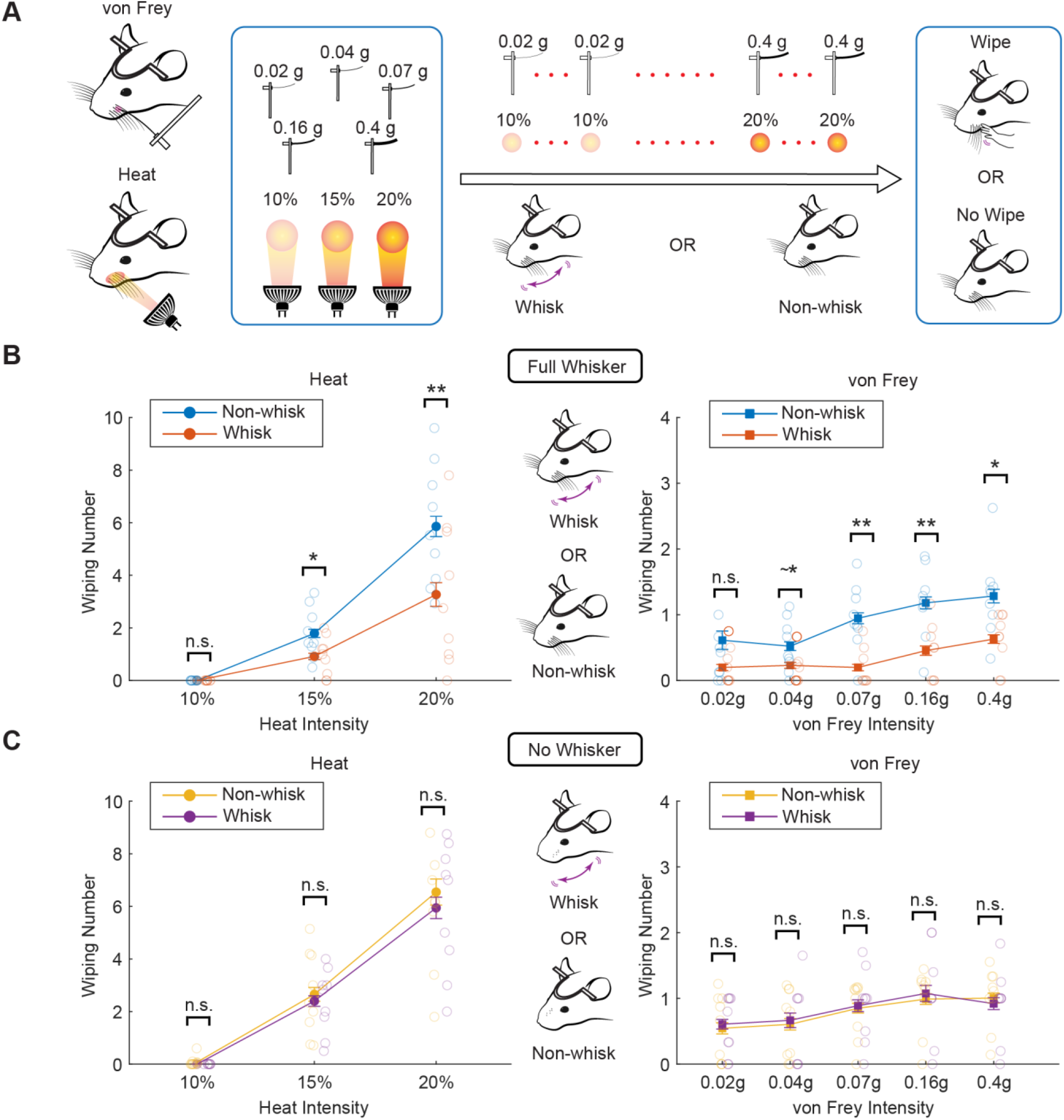
Reafferent signals from self-generated whisking suppressed facial nociception. **A.** Schematic of behavior experiments. Both heat and von Frey stimuli of various intensities were applied to the whisker pad of mice. Mice exhibited either wiping or no-wiping responses, and they were either whisking or resting (no-whisking) during the stimulus delivery. **B.** Nocifensive responses to heat (left) or von Frey (right) stimuli measured by wiping numbers in mice with full whiskers (mean and s.e.m.). The trials were divided into whisking versus no-whisking conditions. **C.** Similar to **B,** except that stimuli were applied after all whiskers were trimmed (no-whisker). (n=9, 5 male and 4 female for B and C). *, p < 0.025; **, p < 0.01; n.s., no significance. Paired t-test was used.

To determine whether this reduction in nocifensive responses to noxious heat/mechanical stimuli was due to vibrotactile sensory signals generated by whisking (i.e., reafference) or due to an efferent signal (i.e., motor command for self-initiated whisking), we trimmed all the whiskers and repeated the same experiments. Even without whiskers, the whisking periods were apparent from facial muscle movements (Movie S1). Importantly, nocifensive face wiping was *not* statistically different between whisking and non-whisking trials when whiskers were removed (Fig. 1C and Fig. S1B). Similar results were obtained when only the whiskers on the same side as the noxious stimulation were trimmed (C57BL/6J, n = 7, with 3 male and 4 female, Fig. S1C). These results indicate that tactile reafferent signals derived from whisking in air, but not motor efference, play a critical role in suppressing facial nociception. Interestingly, after shaving all whiskers (but not with full whiskers, or with only contralateral whiskers), mice appeared to have lost the ability of discriminating different intensities of von Frey stimulation (similar responses to 0.07, 0.16, 0.4g, Fig. 1C). It might be that losing all whiskers on both sides caused some central changes that rendered mice more sensitive to mechanical stimuli (hyper-sensitive), thereby all von Frey stimuli, even those with low force, were perceived as painful non-discriminatively. Nonetheless, with intact whiskers, this behavioral paradigm provides a nice model to study the potential role of S1B (if any) in touch mediated facial analgesia.

### S1B is activated by noxious stimuli

We first briefly summarize the pathways through which tactile and nociceptive signals from whiskers and whisker pad can both reach S1B (Fig. 2A) (*23*). Whiskers and whisker pad skin are innervated by the peripheral axons of trigeminal ganglion (TG) sensory neurons whose central axons project to the principal sensory nucleus (PrV), which receives pure tactile (non-noxious) sensory inputs, and to the trigeminal spinal nucleus (SpV), which receives both innocuous tactile and nociceptive inputs. Tactile signals are relayed from PrV to the contralateral dorsomedial region of the ventral posterior medial thalamus (VPMdm), and then to layer 4 (L4) and layer 5B (L5B) of S1B. Nociceptive information are relayed from SpV to several thalamic nuclei, and to S1B, secondary somatosensory cortex (S2) and insular cortex (IC) (Fig. 2A) (*24–27*). While S2 and IC have long been recognized for their role in processing nociceptive information (temperature and pain), through the extensive intracortical connections between S2 and S1, and between IC and S1/S2 (*28–33*), noxious inputs to S2/IC can also reach S1B, positioning S1B as a site for touch/pain signal interaction.

**Fig.2.**
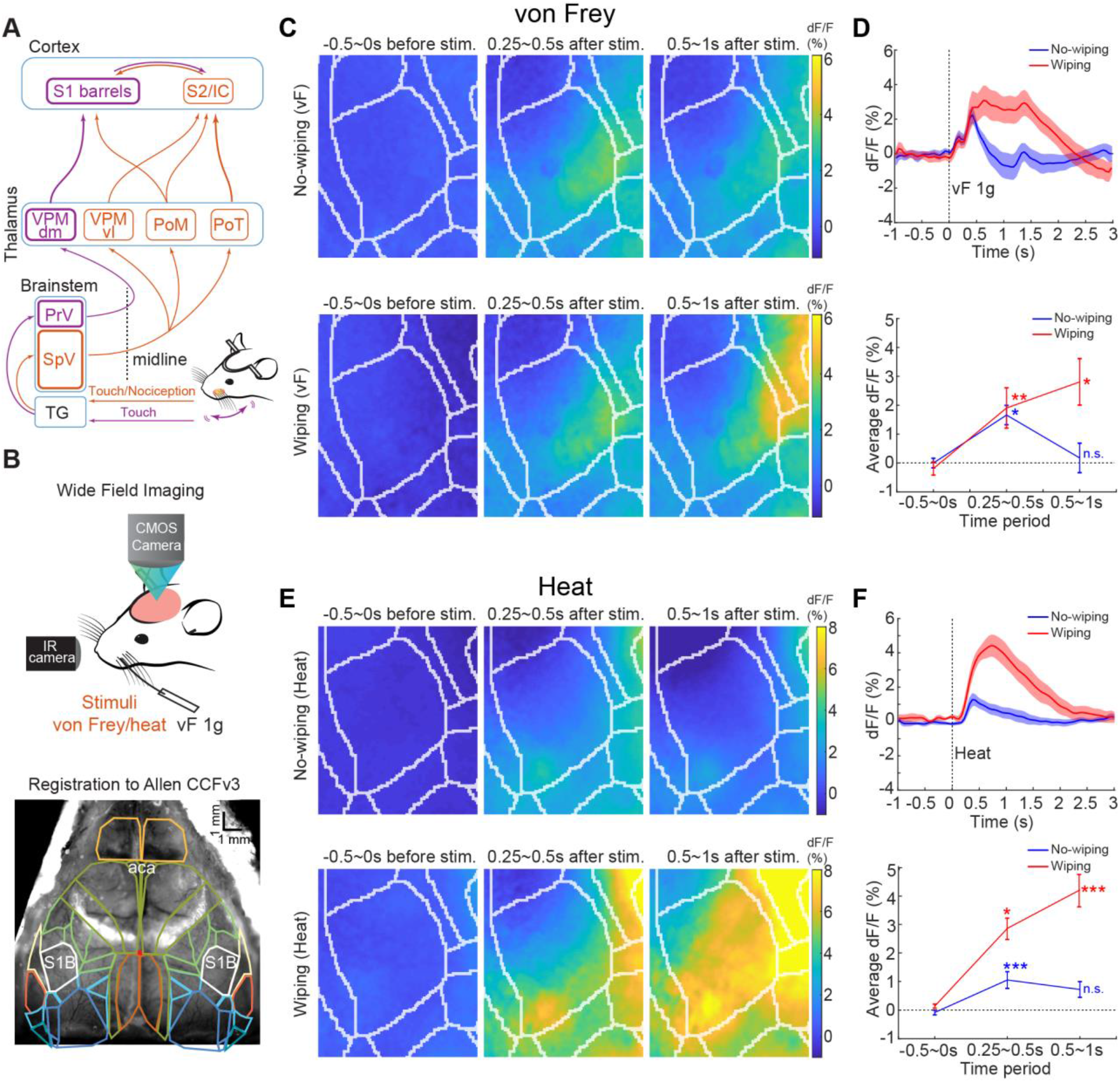
Wide field calcium imaging showing that S1B is activated by noxious stimuli. **A.** Schematic diagrams showing the neuroanatomical pathways relaying touch and noxious signals experienced by the whisker pad from the periphery to different cortical regions. TG, trigeminal ganglion; SpV, trigeminal spinal nucleus; PrV, principal sensory nucleus; VPMdm, dorsomedial region of the ventral posterior medial thalamus; VPMvl, ventrolateral part of VPM; PoM, medial part of the posterior nucleus of the thalamus; PoT, the posterior triangular nucleus of the thalamus; IC, insular cortex. **B**. Top: Setup for wide field imaging of calcium activity. Mice expressing GCaMP7f (C57BL/6J injected retro-orbitally with AAV PHP.eB.syn.jGCaMP7f) are imaged during von Frey stimulation of the whisker pad using a macroscope (see Methods). Bottom: the Allen atlas CCFv3 is scaled and rotated to fit the position of the imaged barrels and to register the borders of the barrel cortex to the surface vessels. **C**. Examples of dF/F maps of the barrel cortex imaged in a mouse stimulated with 1g von Frey fiber. Each map corresponds to the dF/F before stimulation (−0.5s to 0s), when the stimulus first touches the face (0.25 to 0.5s) and during the wiping period (0.5 to 1s). Responses were averaged across trials with (lower panel, n = 16 trials) or without wipes (upper panel, n = 21 trials). **D**. Upper: average response across mice (n = 5) with and without wipes (red: wipes, blue: no wipes, shaded area: +/- sem). Lower: average response during different time periods. The response to von Frey (0.25 to 0.5s after stimulus onset) is significantly greater than baseline during both wipe and no wipe trials. The 2 time courses do not differ during this period but diverge during the wiping period (0.5s to 1s). **E**. Examples of dF/F maps of the barrel cortex imaged in a mouse stimulated with 225ms laser (1450nm) pulse. Each map corresponds to the dF/F before stimulation (−0.5s to 0s), when the stimulus first touches the face (0.25s to 0.5s) and during the wiping period (0.5 to 1s). Responses were averaged across trials with (lower panel, n = 5 trials) or without wipes (upper panel, n = 15 trials). **F**. Upper: average response across mice (n = 6) with and without wipes (red: wipes, blue: no wipes, shaded area: +/- sem). Lower: average response during different time periods. The response to laser (0.25s to 0.5s after stimulus onset) is significantly greater than baseline during both wipes and no wipes trials. The later part of the response (0.5 to 1s) is significantly greater than baseline for trials with wipes but not for trials without wipe. n = 5 in **D** and n = 6 mice in **F**. *, p < 0.025; **, p < 0.01, ***, p < 1×10^-4^, n.s., no significance. Repeated measures ANOVA was used.

To obtain initial evidence that noxious stimuli applied to the whisker pad can activate S1B (directly or indirectly), we performed wide field calcium imaging of the cortical surface (See Methods; Fig. 2B top and Fig. S2A), while applying either infrared laser (wavelength 1450nm) heat or noxious mechanical stimuli using the von Frey filaments to the whisker pad. Mice with cortical-wide GCaMP7f expressions (n = 6 in heat and n = 5 in von Frey experiment) were used for wide field imaging. We first imaged cortical responses to piezo stimulation of individual whiskers to locate the borders of S1B (Fig. 2B bottom and Fig. S2B). Subsequently, we applied laser heat (output power: 300mW, duration ~200-300ms) or von Frey filament stimulations to the whisker pad (1g, which is considered a noxious stimulus since it elicits wiping in more than 85% of trials) and recorded both the behavior and S1B calcium signals (Movie S2). We observed clear S1B activations in response to both noxious laser heat and mechanical stimulation (Fig. 2C-F; referenced to the baseline, von Frey 0.25 ~ 0.5s: p < 0.01 for wiping trials and p < 0.025 for no-wiping trials; von Frey 0.5 ~ 1s: p < 0.025 for wiping trials and p = 0.19 for no-wiping trials; Heat 0.25 ~ 0.5s: p < 0.025 for wiping trials and p < 1 × 10^-4^ for no-wiping trials; Heat 0.5 ~ 1s: p < 1 × 10^-4^ for wiping trials and p = 0.51 for no-wiping trials). In the wiping trials, S1B activity was either prolonged or had a second peak, most likely reflecting prolonged touch signals resulting from wiping of the whisker pad area (Fig. 2D and F). Thus, S1B can indeed be activated by noxious heat/mechanical stimuli applied to facial skin in addition to its well-known role in processing innocuous touch derived from whiskers. Note that motor (M1) and premotor (M2) cortices were also activated, but their signals did not proceed S1 (Fig. S2C), hence S1 activation was not a consequence of M1/M2 activation.

### Ntng1-Cre labels VPM neurons that mainly convey touch but not noxious signals to S1B

The existing anatomical pathways and our wide field imaging results indicate that S1B could process both innocuous and noxious stimuli experienced by the whisker/whisker pad, and thereby potentially be involved in self-whisking mediated facial analgesia. Since VPMdm specifically relays whisker-derived touch information to S1B, we first wanted to know whether blocking this “pure” tactile input pathway to S1B would abolish whisking reafference signal induced suppression of nocifensive responses. We recently generated a Ntng1-Cre knock-in mouse line in which Cre is expressed under the control of the endogenous Ntng1 promoter (*34*). Ntng1, which encodes Netrin-G1 protein, has been shown to be expressed in the dorsal thalamus, and Netrin-G1 likely acts as a guidance molecule for thalamic axons projecting to S1 (*35*). To examine whether Ntng1-Cre expressing VPM neurons (VPM_Ntng1_) relay tactile information to S1B, we first injected the AAV-Flex-GFP unilaterally into VPM of Ntng1-Cre mice and confirmed anatomically that the GFP labeled VPM_Ntng1_ neurons send dense axon projections mainly into the ipsilateral S1B to L4 and L5B (Fig. 3A upper). Next, to examine the in vivo activity of VPM_Ntng1_ neurons, we injected AAV-Flex-GCaMP6m in the VPM of the Ntng1-Cre mice (Fig. 3A bottom), and performed fiber photometry recording of their responses in different behavioral conditions.

**Fig. 3.**
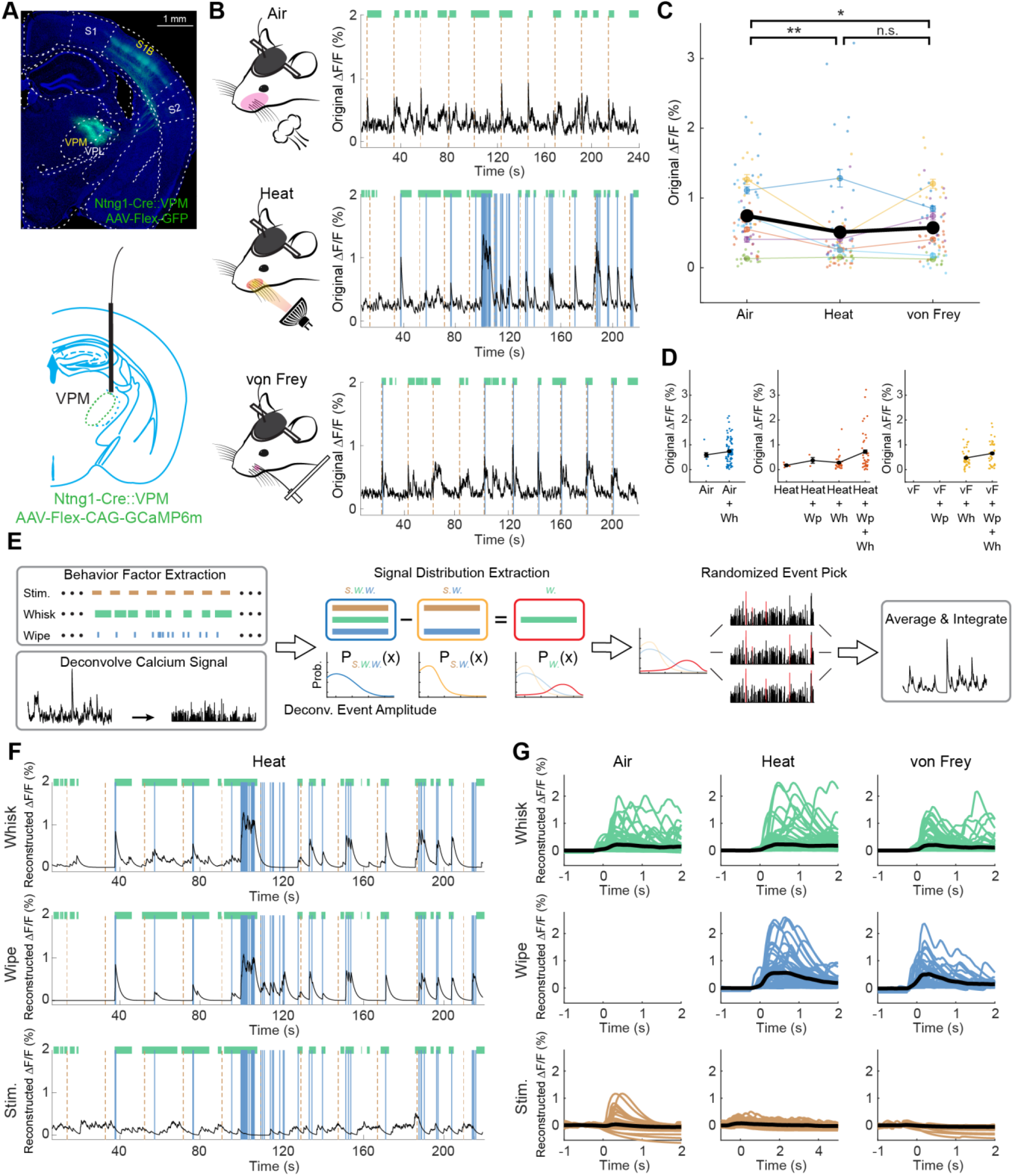
VPM_Ntng1_ neurons mainly convey touch but not noxious signals to S1B. **A.** Upper panel, representative image of GFP labeled VPM_Ntng1_ neurons, and their axonal projections in the ipsilateral S1B layer 4 and 5B. Bottom panel, schematic for the fiber photometry recording of the population activity from the VPM_Ntng1-GCaMP6m_ neurons. **B.** Representative VPM_Ntng1_ population activity traces recorded by fiber photometry in different conditions. Air puff, heat and von Frey stimuli were applied to the whisker pad of mice. Green bars, whisking periods. Orange dotted lines, stimulus onsets. Blue solid lines, wiping moments. **C.** Maximum intensity of raw dF/F_0_ from all trials for different stimulus types. Different colored dots represent trials from the same mice (n = 6 different mice). The colored line charts are the averages of maximum intensity of individual mice, while the black line chart is the grand average. **D.** Maximum intensity of raw dF/F_0_ from all trials separated further based on factors (stimulus, whisking and wiping). **E.** Schematics of the factor-related signal estimation algorithm with binary factor labeling. **F.** Example traces of factor separated traces from one session of heat stimuli trials. **G.** Separated signals epoched based on factors which were pre-factor corrected and aligned to the factor onset (t = 0) in different conditions, each line represents a trail from a mouse and results from all mice are shown. The thick black line represents the grand average. *, p < 0.025, **, p < 0.01, n.s., no significance. Two sample t-test was used.

Using the same head-fixed behavior setup described above, we applied either innocuous stimulus (air puff) or the two noxious stimuli (noxious heat and von Frey mechanical stimuli) onto the whisker pad, while recording the population calcium signal from the VPM_Ntng1_ neurons (Fig. 3B, n = 6). The behavior was simultaneously filmed and the periods of whisking and wiping events were manually tracked. We first compared the maximum intensity of dF/F_0_ from each trial for different stimulus types (Fig. 3C; see Methods). On average, the VPM_Ntng1_ neurons responded to air puff with stronger signal intensities than to heat or to von Frey stimuli, consistent with the primary role of VPM in signaling innocuous touch (Fig. 3C, *p* < 0.01 to heat, *p* < 0.025 to von Frey), and there was no statistical difference between responses to the two different noxious stimuli (Fig. 3C, *p* = 0.485). Because in each session, mice sometimes whisked when stimuli were applied or wiped their faces in response to the stimuli (note that wiping was never induced by air puff), we further separated the trials based on these two behavioral factors and then compared the maximum signal intensity. Whisking and wiping both increased the population activity when compared to the no-whisking or no-wiping conditions in responses to the same stimulus type (Fig. 3D).

Due to the slow decay of GCaMP signals, the raw dF/F_0_ population activity signals represent a mixture of external stimulus-elicited, as well as self-initiated whisking and wiping generated signals. We therefore asked what the “pure” signals corresponding to different stimuli (air puff, heat, von Frey), or to different behaviors (whisking or wiping) were for VPM_Ntng1_ neurons. We first deconvolved the dF/F_0_ signals into individual population calcium events using a constrained foopsi method (*36*), and then applied a signal separation algorithm that would extract the traces related only to one factor of interest (factor = stimulus, whisking or wiping) (Fig. 3E; see Methods). Using heat trials as an example (Fig. 3F), this algorithm separated the raw dF/F_0_ trace into “pure” whisk-, wipe-, and heat-related signals. In this case, the signals induced by whisking and wiping (both behaviors generate reafferent touch signals from whisker pad) were markedly higher than induced by heat. We performed this analysis for all trial types and the pooled results are shown in Fig. 3G. As expected, the activity of VPM_Ntng1_ neurons was dominated by tactile signals from whisking, wiping, and air puff, and was minimally responsive to noxious heat or mechanical stimuli (Fig. 3G). These results indicate that in awake behaving mice, VPM_Ntng1_ neurons indeed predominantly relay vibrotactile touch signals into the S1B.

### Inhibiting activity of S1B-projecting VPM_Ntng1_ neurons diminished whisking induced suppression of nocifensive responses

Next, we investigated whether blocking the main source of whisker-derived touch signals into S1B by inhibiting VPM_Ntng1_ neurons would abolish the whisking induced suppression of nocifensive responses. We injected AAV-Flex-PSAM^L141F^-GlyR-GFP into the VPM of Ntng1-Cre mice for transiently chemogenetic silencing VPM_Ntng1_ neurons (Fig. 4A, left). PSEM308, the PSAM agonist, opens the PSAM^L141F^-GlyR chimeric ion channels to allow influx of chloride ions, thereby inhibiting the activity of the neurons expressing PSAM^L141F^-GlyR (*11, 37*). Animals’ responses to heat and von Frey stimuli (Fig. 4A, right) were measured at baseline (both pre- and post-PSEM sessions) and 20 min after PSEM308 administration (Tocris, 5 mg/kg, i.p.).

**Fig. 4.**
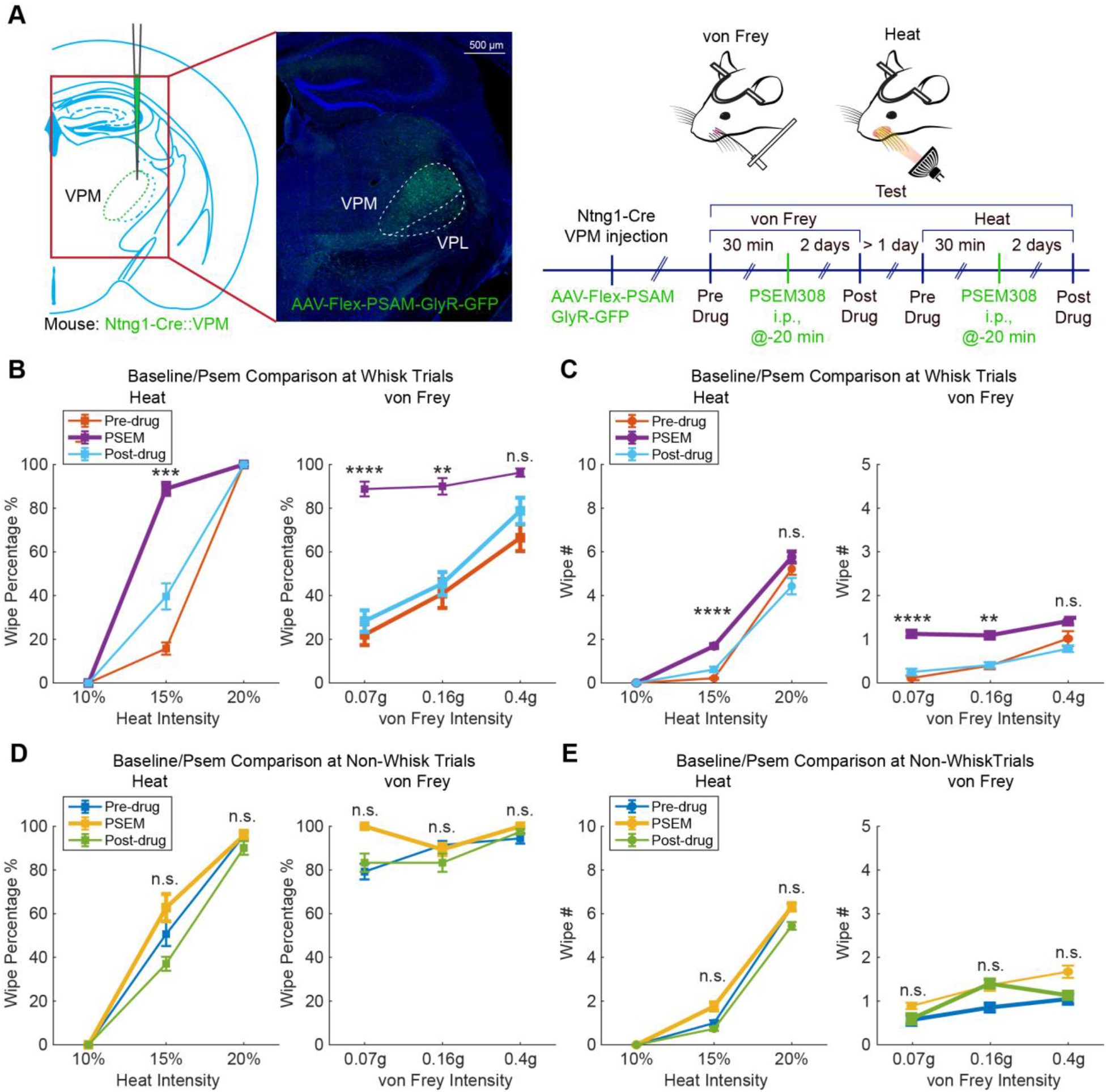
Inhibiting activity of VPM_Ntng1_-S1B projecting neurons diminished whisking induced suppression of nocifensive responses. **A.** Left: schematic showing the manipulation strategy by injecting AAV-Flex-PSAM-GlyR-GFP into the VPM of Ntng1-Cre mice, and a representative image of the post hoc histology. Right: timeline of von Frey and heat tests to assess the effect of the inhibition of VPM_Ntng1_ neurons on the behavior. **B-C.** Nocifensive behaviors in whisking trials of pre-drug, PSEM administered, and post-drug sessions, measured by percentage of trials with wiping responses (**B**) or wiping numbers (**C**). **D-E.** Nocifensive behaviors in no-whisking trials of pre-drug, PSEM administered, and post-drug sessions, measured by percentage of wiping trials (**D**) or wiping numbers (**E**). n = 8 mice. *, p < 0.025, **, p < 0.01, ***, p < 1×10^-4^, ****, p < 1×10^-6^, n.s., no significance. Repeated measures ANOVA with post-hoc Tukey’s honestly significant difference was used. Linear mixed-effect model was further applied to the highest intensities for the wipe number measurement in both heat and von Frey experiments.

Again, we separated trials into whisking versus non-whisking ones. Analysis of trials with self-generated whisking revealed that during PSEM mediated inhibition of VPM_Ntng1_ neurons, both the percentage of trials with face wiping and the wiping numbers increased significantly in response to noxious heat and von Frey stimuli compared to those of no drug conditions (pre- and post-drug baselines; Fig. 4B, p < 0.01, 1 × 10^-4^ or 1 × 10^-6^; Fig. 4C, p < 0.01 or 1 × 10^-6^; n = 8). In other words, the whisking mediated anti-nociception disappeared upon VPM_Ntng1_ inhibition. As controls, PSEM treatment did not alter wiping responses during non-whisking periods, i.e., wiping behavior remained at the similar levels during the non-whisking trials in PSEM sessions compared to the baselines (Fig. 4D and E. p > 0.1 using repeated measures ANOVA). Thus, inhibiting VPM_Ntng1_ neurons did not alter normal nociception, consistent with our imaging results that these neurons do not convey nociceptive signals as they are “pure” tactile responders. Noxious information is relayed to the cortex by other thalamic and non-thalamic neurons. Together, these results indicate that whisking mediated analgesia is diminished if the reafferent tactile signals are blocked from transmitting to S1B through the canonical VPM-S1 pathway. Since the main axonal target of VPM_Ntng1_ neurons is S1B, these findings further underscore the essential role of S1B in integrating tactile and painful information to enable touch mediated analgesia.

### In vivo miniscope imaging of S1B neuronal responses to noxious stimuli in behaving mice

Touch information transmitted by VPM_Ntng1_ neurons to L4 S1B neurons is further relayed to L2/3 neurons. Nociceptive information could also arrive at upper layers of S1B either through the PoM pathway or via the inter-cortical connections with S2 and IC (Fig. 2A). Thus, L2/3 of S1B could be a node where tactile and noxious signals interact. To investigate how S1B might be involved in the touch induced reduction in pain responses, we asked two main questions: first, how S1B differentiates innocuous versus noxious stimuli at the individual neuron level, and second, how S1B neural population represents whisking induced suppression of nociception. To answer these questions, we decided to use in vivo imaging approaches to characterize S1B L2/3 activity. To ensure consistent imaging of the same S1B region across mice, we used intrinsic signal optical imaging to locate the activation area of the C2 whisker barrel in S1 cortex (Fig. S3, see Methods), and then injected AAV-Syn-GCaMP6f into the layer 2/3 of the C2 barrel area in all animals (n = 6). We implanted the GRIN lens above the injection site to image calcium responses of individual S1B neurons using microendoscope/miniscope (Inscopix).

We performed in vivo imaging in head-fixed mice running on the wheel while applying innocuous (air puff) or noxious stimuli (heat, 15% intensity; von Frey, 0.4 g) to the whisker pad (n = 6; 6 sessions per mouse, see Methods). Calcium traces (dF/F_0_) and the spatial footprints of the individual recorded ROIs were extracted using MIN1PIPE (*38*) (Fig. 5A and B). Pooling the trial average of all the neurons (n = 764 and 782 neurons, respectively, for heat and von Frey experiments from 6 mice) and then sorting neurons by their peak activity following the onset of either heat or von Frey stimulus delivery revealed that different neurons were activated at different timepoints in the 10 seconds window following stimulus onset, in both whisking and non-whisking trial types (Fig. 5C). We also marked the time of wiping events (blue dots above the heat maps in Fig. 5C), to show that in general, it took mice 3-4 seconds to wipe their face in response to heat (likely due to the slow heating up of the tissue, and/or slower conduction velocity of heat-responsive c-fibers); whereas they wiped within 1 second upon von Frey stimulation (as noxious mechanical stimuli likely activate the fast conducting A*δ* sensory fibers). In both heat and von Frey trials, there were subgroups of neurons with the rising phase of their calcium activity peaked prior to the onset of wiping behavior, suggesting that these subsets of neurons may signal the presence of noxious stimuli (Fig. 5C; see Fig. S4 for alternative sorting strategies showing the nociceptive signal component and the dominant touch/wiping signal component). However again, due to the intermingled nature of the signals likely reflecting a combination of external stimuli, whisking and wiping behaviors, it is difficult to draw conclusions based on the trial averaged signals.

**Fig. 5.**
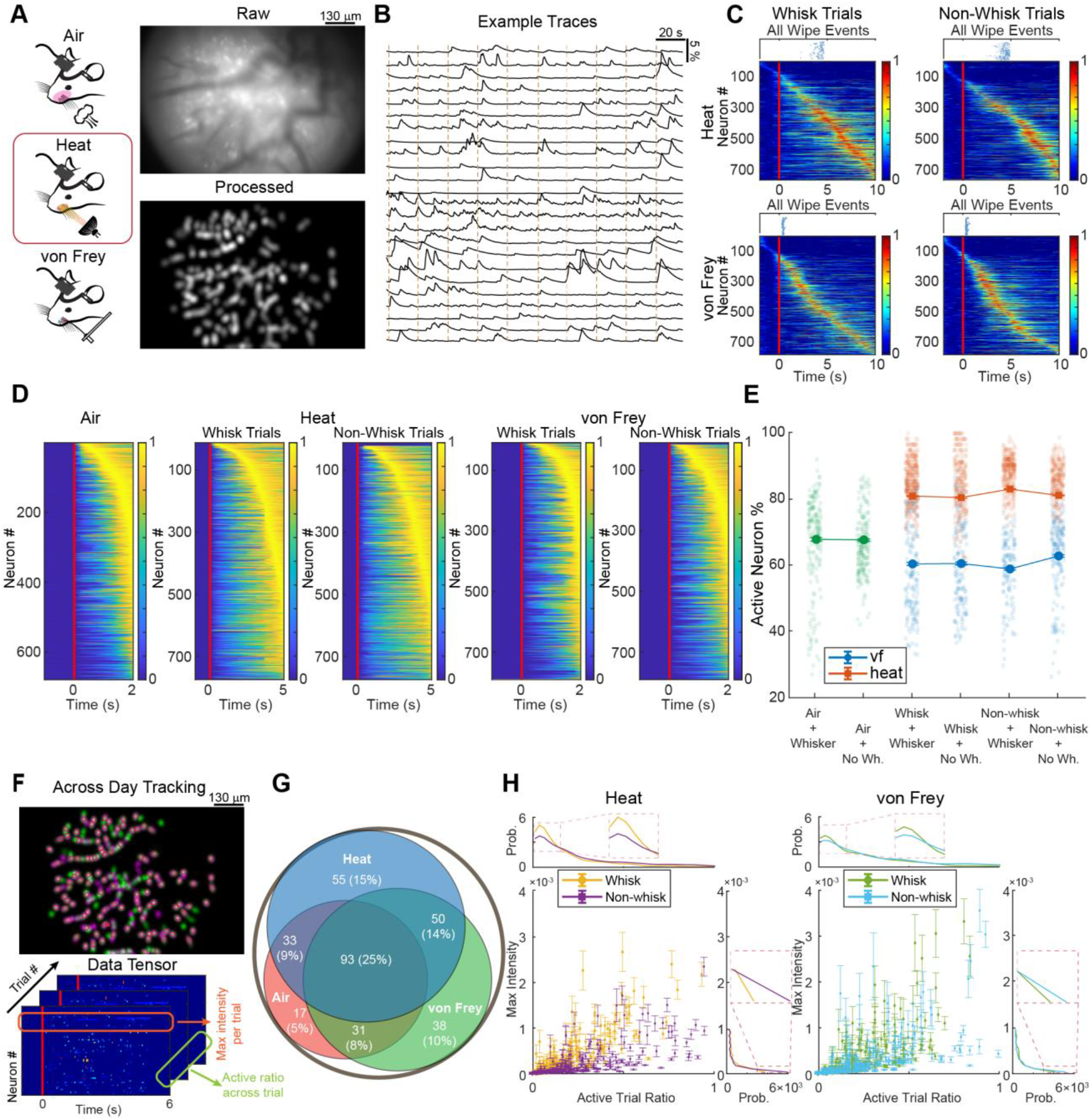
S1B L2/3 contains modality-specific and non-selective neurons and whisking reduces their active trial ratios. **A.** Left: in vivo imaging while applying innocuous or noxious stimulus. Right: raw field of view (top) and processed ROIs (bottom) of one representative imaging session. **B.** Example traces of the extracted calcium fluorescence from the representative ROIs in an imaging session. Orange dashed lines indicating stimulus (heat) onsets. **C.** Trial average of raw dF/F_0_ for each neuron aligned to the stimulus onset (red lines). The traces were normalized. The first wiping events of all trials were also concatenated vertically and shown as blue dots on top of each heatmap. **D.** Trial average of stimulus specific signals (rescaled to [0, 1]) for each neuron aligned to the stimulus onset (red lines). **E.** The percentage of active neurons per trial in response to different stimulus types (green, air puff; red, heat; blue, von Frey). The trials for noxious stimuli were further separated into whisking versus no-whisking, whisker versus no-whisker conditions. **F.** Top: an example image of the same-neuron tracking for two different sessions of the same mouse. Red circles, tracked same neurons. Bottom: a demonstration of how to extract the two measures, maximum intensity per trial and active trial percentage, from the data tensor. **G.** Venn diagram showing the distribution of stimulus activated neurons (both absolute numbers and percentage of all the tracked neurons). **H.** Scatter plots with mean and s.e.m. as a function of the maximum intensity per trial and active trial percentage, and the marginal distribution as a function of each variable. The trials were separated into whisking and no-whisking conditions.

Therefore, we adapted the signal separation algorithm used above for the univariate fiber photometry data to the multivariate (i.e., multiple ROIs) miniscope calcium imaging data. We focused on isolating the specific stimulus-related signals to examine whether S1B activity can distinguish different types of stimuli (innocuous air puff, noxious thermal, or noxious mechanical stimuli). For different stimuli, we used different time windows following the onset of stimuli. Since we used an air blaster to deliver air puff, the stimulus lasted about 1-2 seconds. Considering the slower kinetics of GCaMP, we therefore analyzed imaging data up to 2 seconds following the onset of air puffs. For heat stimulations, the heating lamp was automatically turned off after 5 seconds and mice showed wiping at around 3-4 seconds, thus, we analyzed signals up to 5 seconds. For von Frey, we manually applied von Frey for less than 1 second then retracted the fiber, and wiping occurred at around 1 second (up to 2 seconds; wiping pushed away the von Frey filament thereby terminating the mechanical stimulus sooner), thus we analyzed calcium signals up to 2 seconds. The extracted stimulus-specific signals for imaged neurons were plotted as trial averaged traces and sorted by their peak timing (Fig. 5D). These analyses revealed that S1B L2/3 neurons indeed respond to both innocuous air puff to whiskers (as expected) and to noxious heat and mechanical stimuli applied to the whisker pad (with time course correlating with the duration of the stimuli), in both whisking and non-whisking trials (Fig. 5D). Note that the same method was applied to the VPM fiber photometry data where very little heat or von Frey related signals were found (Fig. 3G). Taken together, the results contrast the different roles of VPM_Ntng1_ and S1B L2/3 neurons in touch induced analgesia, with S1B being the potential site integrating touch/pain information. To further validate that we indeed extracted stimulus-specific signals, we applied our method to the baseline periods of imaging (where the data were not labeled with any of the factors of interest, namely stimulus, whisking or wiping), with the random labeling (i.e., assigned pseudo stimulus, whisking or wiping events). We then extracted the pseudo factor related signals and calculated the trial average for all neurons and did not find any obvious calcium events corresponding to pseudo stimuli (Fig. S5).

We further examined whether S1B population activity patterns showed modality-dependent (gentle touch, heat, noxious touch) differences. For each trial, we used the stimulus-specific deconvolved calcium signals to calculate the adaptive threshold for each neuron, and then extracted the number of active neurons (i.e. neurons with activity above the thresholds; defined by having at least 1 calcium event larger than 1 standard deviation of the signal in each trial, with trial duration for air puff = 2 seconds, heat = 5 seconds, and von Frey = 2 seconds) (see Methods). We then pooled the data from the same types of trials and plotted the distribution of the active neuron ratios for each type (active neurons / all neurons per session; Fig. 5E). The results showed that on average, ~67% of imaged S1B L2/3 neurons had calcium events in air puff trials, whereas ~80% or 60% of imaged neurons were active in noxious heat and von Frey trials, respectively (Fig. 5E). The stimulus-dependent active neurons ratios were similar in whisking/non-whisking, and in full whisker/no whisker conditions. The differences in percentages of active neurons likely reflected the different stimulus duration (1-2 seconds for air puff, 4-5 seconds for heat, and 1 second for von Frey), as well as the size of the area affected by the stimulus (full whisker pad for air and heat, versus a small locus on whisker pad for von Frey). The results suggest that the ratios of active neurons could be used to discriminate stimulus type.

To further determine whether S1B L2/3 contains neurons that are specifically tuned to different stimulus modalities, we tracked neurons imaged in each mouse across different sessions, using the modified CellReg algorithm (*39*), and examined their responses to different stimuli (Fig. 5F). Of all the neurons, we successfully tracked 365 neurons. Using the same adaptive thresholding approach (see Methods), we determined whether each tracked neuron was activated in response to air puff, heat or von Frey (Fig. 5G). The Venn diagram shows that although a subgroup of neurons (n = 93) non-specifically responded to all three types of stimuli, there were comparable subgroups of neurons that were either active in two types of stimuli or only specifically activated by a single type of stimulus (Fig. 5G). Thus, in principle, the modality-specific single neurons, together with the overall ratios of active neurons mentioned above, could enable S1B to discriminate touch versus pain and also the types of noxious stimuli.

### Analyzing the effect of whisking on activity of individual S1B L2/3 neurons in response to noxious stimuli

While the analyses described above provide evidence for the involvement of S1B in processing and potentially discriminating gentle versus noxious touch versus heat, it did not reveal how whisking-derived reafference touch signals might alter how S1B neurons process noxious information. To begin to understand this question, in the same-cell tracking process, we also extracted the maximum intensity per neuron per trial, and the percentage of active trials for each tracked neuron (active trials / all trials per session) using the deconvolved stimulus-specific calcium signals (active trials defined by having maximum intensity of deconvolved events above one standard deviation of all trials’ maximum intensity, see Methods for details, Fig. 5F). We further separated trial types into whisking versus non-whisking ones, as shown in Fig. 5H, main panels. We also calculated the probability density function for these two measures (Fig. 5H, top and right panels).

We found that whisking slightly decreased the proportion of neurons with lowest maximum intensity (see probability density on the right side of the heat and von Frey plots in Fig. 5H; Kolmogorov-Smirnov test (KS test) with p < 0.001 for heat but p > 0.01 for von Frey using trial averaged data; however, if using all trial data rather than trial-averaged data, the KS test showed significance with p < 0.001 for both conditions). In other words, whisking increased the baseline activity of those inactive/silent neurons. This is likely due to the propagation of whisking signals in S1B that increased basal activity of L2/3 neurons. However, the probability density at high maximum intensity was almost unaltered by whisking. On the other hand, our analyses also revealed that the majority of S1B L2/3 neurons were only activated (compared to their own baseline activity) in low percentages of trials and had low activity levels, consistent with the sparse coding mode of L2/3 neurons observed previously (*40*). Importantly, whisking further reduced S1B neurons’ overall responsiveness to heat and von Frey by increasing the proportions of neurons with low percentage of active trials (see probability density in top panels of Fig. 5H; KS test showing p < 0.01 for heat but p > 0.01 for von Frey using trial averaged data; however, if using all trial data rather than trial-averaged data, the KS test showed significance with p < 0.001 for both conditions). In other words, whisking makes S1B L2/3 neurons as a population less likely to respond to subsequent noxious stimuli. This result is consistent with previous studies revealing that whisker-derived inputs to L4 can mediate feedforward inhibition of L2/3 neurons (*18, 24*). While previous work only examined tactile processing, our results suggest that this whisking-derived feedforward inhibitory mechanism also contributes to the suppression of S1B nociceptive responses.

### Tensor component analysis revealed the differences in S1B population activity between trials with or without nocifensive responses

To further investigate what aspects of S1B population activity patterns (rather than highest intensity) correlate with nociception, we performed tensor component analysis (TCA) (*41*) to analyze the raw dF/F_0_ data in heat and von Frey trials. TCA, compared to the traditional trial average principal component analysis (PCA), enables dimension reduction capturing single-trial dynamics. Briefly using the method described by (*41*) as shown in Fig. 6A, we applied TCA to decompose the data tensors (a collection of matrices) obtained from raw fluorescence signals into factors corresponding to neuronal (i.e., assemblies of neurons), temporal (i.e., neural dynamics shared by all trials), and trial (i.e., trial-to-trial scale changes of the same temporal factor) components (Fig. 6A; Fig. S6; see Methods). Because we used wiping behavior as the surrogate for pain perception, we mainly wanted to identify the differences in neuronal patterns between wipe and no-wipe conditions in both heat and von Frey trials. We extracted multiple tensor components (TCs) per mouse for each type of trial whose trial factors had the highest similarity score with the actual behavior readouts (wipe versus no wipe, and further separated into whisking versus no whisking trials) (Fig. 6B).

**Fig. 6.**
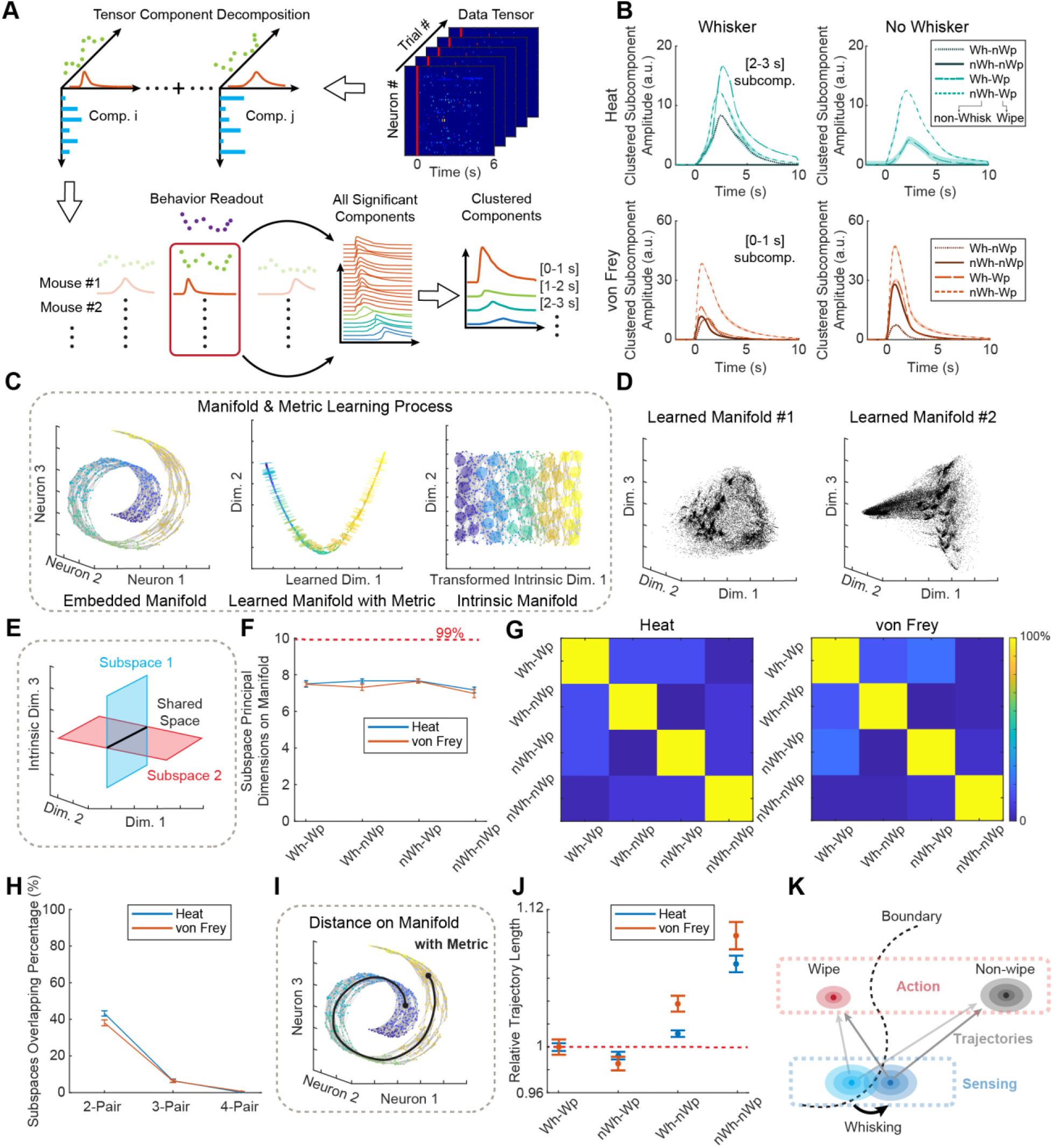
Whisking moves the S1B population dynamics towards the no-nociception-no-action states on the intrinsic manifold. **A.** Schematic of the processing pipeline using TCA. After TCs were extracted, we pooled the ones in the same 1-second time bin into “clustered components” and compared these between wiping and no-wiping trials. **B.** The amplitudes of clustered components for different behavioral combinations (whisking/no whisking, wiping/no-wiping) in response to heat or von Frey stimulation in full whisker and no whisker conditions. The no-wiping conditions in the heat experiments did not have TCs in the time bin (zero amplitude). **C.** Steps to learn the intrinsic manifold and to estimate the point-wise metric, shown with a toy model (swiss roll). See methods. **D.** Examples of learned manifolds (without correction with metric) from two sessions. The states (individual black dots) were projected onto the first three dimensions of the intrinsic dimensions learned with LEM. **E.** Schematics for how we define shared space by the two subspaces on the manifold formed by the two different behavioral configurations. **F.** The average number of dimensions of S1B neural activity in each behavioral configuration from PCA. The first n principal components (dimensions) whose cumulative variance of neural activity explained exceeded 99% were preserved. The red dotted line indicates the dimension that has 99% of variance in an isotropic space. **G.** The numbers of dimension of the shared space between any pair of subspaces formed by different behavioral configurations, shown in the form of heatmap matrix. Diagonal elements indicate the shared space by each subspace itself, which are always 100%. **H.** The general summary of the dimensions of the shared space across any combination of the subspaces formed by different behavioral configurations. **I.** Schematic for how trajectory length is calculated along the intrinsic manifold. **J.** The summary of the trajectory lengths (mean with s.e.m, averaged across trials) along the manifold, grouped by the behavioral configurations and normalized to the average trajectory length of the whisking-wiping configuration. **K.** Schematic summary S1B population dynamics underlying whisking induced analgesia.

For the extracted TCs from all mice, we placed them into one-second bins (the whole duration of a trial is 5 seconds), and pooled (summed up) the TCs in the same time bin into “clustered components” (illustrated in Fig. 6A). We then compared the amplitudes of the clustered components in the time window right before the wiping behavior (the surrogate for nociception), i.e., [2, 3s] for heat and [0, 1s] for von Frey trials. This analysis showed that in conditions where there was no nocifensive wiping, regardless of whether whiskers were present or all-removed and whether mice were whisking or not whisking, the clustered TCs in the pre-wiping window had smaller amplitude than the ones exhibiting wiping behavior (Fig. 6B, note that in heat experiments, the no-wiping conditions had zero amplitude, thus not observable on the figure). In other words, S1B L2/3 population activity differed between trials with or without nocifensive behaviors in the pre-action window, which could be considered as the window when noxious information might be perceived. Larger clustered components (i.e., total behavior-specific neural dynamics) correlated with subsequent nocifensive actions, consistent with the idea of S1B as a sensorimotor cortex (see Discussion). On the other hand, we attempted to compare wiping trials with or without whisking to uncover the potential role of whisking in suppressing wiping. However, due to the dominance of wiping signals, we did not detect any specific component that could explain how might whisking work in non-wiping trials compared to wiping trials (data not shown).

### Whisking suppresses nocifensive behavior through moving S1B neuronal population dynamics towards the non-painful states on the intrinsic manifold

Since the TCA analysis did not uncover the exact effect of whisking on nociception, we turned to examine the state space formed by the population activity of S1B neurons for each session (i.e. imaged neurons, their activity recorded over time throughout the entire session) to answer this question. All states for a given session in the high dimensional state space (number of dimensions = number of neurons) are assumed to belong to a (continuous) manifold, which has lower intrinsic dimensions embedded in that state space (Fig. S7). To find/approximate the presumed intrinsic manifold, we applied a manifold learning algorithm (*42, 43*), and then calculated the metric that geometrically defines the manifold to reconstruct the Riemannian manifold (Fig. 6C; see Methods)(*44, 45*). Examples of the learned manifolds for two different sessions projected onto three dimensional space are shown in Fig. 6D, where each dot is the projected state from the manifold to the space spanned by the first three dimensions. Using persistent homology analysis (*46*), we found that all learned manifolds were continuous (β0) with no general holes (β1; Fig. S8), suggesting that these manifolds all have the same trivial topological property (single connected component).

Having computed the lower dimensional manifolds, we asked whether the subspaces for different behavioral configurations (i.e., different combinations of whisking/no-whisking and wiping/no-wiping) corresponded to orthogonal dimensions, or largely shared the same subspace (Fig. 6E). In other words, we looked for extent of commonality between population dynamics for different behaviors. In order to answer this question, we further reduced the dimensions of each subspace using PCA and kept the first N components whose cumulative explained variance exceeded 99%. The average number of dimensions stayed relatively constant across all configuration subspaces (Fig. 6F; see Methods). Analysis of the shared dimensions across different subspaces within the same session and mouse showed that most of the dimensions between any two pairs of behavioral configurations were orthogonal in both heat and von Frey trials (Fig. 6G). A general summary of the shared dimensions across all degrees of behavioral configurations further confirmed this observation (Fig. 6H). That subspaces occupied by different types of behaviors are mostly orthogonal suggests that whisking alters S1B population neural dynamics in response to heat or von Frey differently than non-whisking conditions, thereby resulting in different nociception and responses (wiping versus no wiping). It is also possible that in trials with no whisking, other factors (e.g., animal’s attention, internal state, etc.) may alter S1B dynamics to cause lack of nociception (and thus no wiping).

To better define the role of whisking, we measured the length of the trajectory traveled by the activity of S1B neurons on the manifold for each trial (which starts with stimulus onset and ends roughly 5 seconds or 1 second after applying heat or von Frey, respectively). We applied a shortest path algorithm on every two neighboring states of a trial on the learned Riemannian manifolds and summed up the total geodesic distances (Fig. 6I). Since the manifolds do not necessarily scale in geometrically equivalent manner, we normalized the calculated trial trajectory length to the ones with both whisking and wiping (Fig. 6J). In general, the trials with wiping (indicative of nociceptive perception) had shorter trajectories than the trials without wiping in response to both heat and von Frey (Fig. 6J). This is likely due to the fact that nocifensive face wiping upon sensing noxious stimuli is a stereotypical response, with low energy barrier between the state of sensation (nociceptive stimuli sensed) to the state of action (wiping). Notably, among the wiping trials (in which pain is presumably sensed), the trajectory in non-whisking conditions was the shortest, shorter (below the red dashed line in Fig. 6J) than that of wiping trials with whisking. This finding suggests that when S1B neural activity likely signals the detection of noxious stimuli, whisking pulls the trajectory away from nociception thereby lengthening the distance of the neuronal state to the action outcome of wiping. By contrast, in the non-wiping trials (in which the noxious stimuli failed to be sensed), the presence of whisking resulted in a shorter trajectory than that of no-whisking trials. This result suggests that whisking facilitates (i.e. shortens) the trajectory toward the action outcome of non-wiping (Fig. 6K schematic). Thus, our results, together with other studies on the role of S1 in decision making (*47, 48*), support the role S1B L2/3 neurons’ population dynamics in integrating peripheral derived diverse somatosensory stimuli and steering the brain states toward different actions. S1B is an important node in mediating the phenomenon of pain suppression by vibrotactile touch. This is achieved by a mechanism in which self-generated whisking inputs move the initial population dynamics in response to noxious inputs towards the states signaling innocuous touch and thus reducing the need for nocifensive actions.

## Discussion

In this study, we developed a mouse behavioral model of vibrotactile touch mediated suppression of nociception. We showed that reafferent signals from self-generated whisking reduce nocifensive wiping responses to noxious heat and mechanical stimuli applied to the face. This effect requires the transmission of tactile signals through the VPM to S1B pathway, pointing to the involvement of S1B in whisking induced face analgesia. In vivo calcium imaging of S1B L2/3 neurons during this behavioral paradigm revealed the existence of modality-specific and modality-general S1B neurons, and that whisking moderately reduced the overall responsiveness of individual S1B neurons to noxious stimuli. Tensor component analysis further uncovered consistent differences in S1B population activity prior to the different action outcomes (wiping or no-wiping) in response to noxious stimuli, supporting the role of S1B in sensation and sensation-guided-action. Finally, by analyzing the intrinsic manifold equipped with metric, we discovered that S1B activity in wiping (nociception) versus no-wiping (lack of nociception) trials occupy different sub-space on the manifold of S1B state space, and importantly, pre-existing whisking pulls the S1B dynamics away from the nociception-to-wiping trajectory, while facilitates the state transitions along the no-nociception-no-wiping trajectory.

While pain relief by touch is a common human experience, a simple and robust animal behavior model has not been previously established. Instead, electrophysiological studies had largely focused on spinal mechanisms using anesthetized animals. In fact, most animal behavior studies have focused on the opposite phenomenon: pain-induced tactile allodynia (i.e. under painful conditions, animals show hypersensitivity to gentle touch), due to the ease of applying external tactile stimuli to painful areas. While animals clearly use behaviors involving self-generated touch such as licking and wiping of a hurt body part to relieve pain, these behaviors contain mixed components of pain, touch, and movements, and lack a means to measure/report the changes in the perceived pain induced by licking/wiping. Without an easy and reproducible awake behaving animal model, it is difficult to examine whether supraspinal centers (and where) are involved in touch mediated analgesia. Here, taking advantage of the natural whisking behavior in mice, which is readily exhibited in head-fixed preparation, we showed that the whisking-derived vibrotactile reafferent signals can significantly reduce nociception to noxious heat or mechanical stimuli applied to the face, thereby establishing a simple yet robust behavioral paradigm for studying touch induced analgesia. Note that we used face-wiping as a surrogate for pain perception by the mouse. Unlike the simple head withdrawing away from the stimuli (in head-free mice), this goal-directed action of moving the ipsilateral forelimb and forepaw toward the face and wiping the stimulated region is not a simple low-level reflex. This behavior likely requires localizing the stimulus to the face, perceiving the stimulus as noxious, and generating a motor action to relieve the transient pain, making it a reasonable surrogate for pain perception.

Using this natural behavior paradigm, we uncovered the role of S1B barrel cortex in whisking mediated face analgesia, since blocking the tactile specific thalamocortical pathway from VPM_Ntng1_ to S1B largely abolished the analgesic effect of whisking, but had no effect on nociceptive responses in non-whisking conditions. A long history of studies on the rodent S1B has highlighted its role as the primary cortical region that processes whisker-derived tactile information (*16, 17*). The well-known functions of S1B include object localization and texture perception, as well as integration touch with locomotion. However, whether and how S1B is involved in processing painful stimuli experienced by the face (whisker skin) have been poorly studied especially in awake behaving rodents. Part of the difficulties of studying pain processing in S1B in awake animals is that noxious stimuli elicit immediate nocifensive behaviors, making it hard to separate stimulus-related versus behavior/movement-related signals. The slower decay of the calcium indicator makes separating signals even more of an issue. In fact, recent brain-wide electrophysiological recordings or calcium imaging studies all revealed that behavior/action signals are prominent and widely spread throughout the brain (*49–51*). Here we developed a signal estimation algorithm that enables extraction of specific neural signals corresponding to stimulus or behavior from both fiber-photometry and calcium imaging data. Using this algorithm, we confirmed that VPM_Ntng1_ neurons relay primarily tactile stimuli but are minimally responsive to nociceptive stimuli. Furthermore, we found that S1B L2/3 neurons can indeed be activated by innocuous touch, heat, and noxious mechanical stimuli. Our wide-field imaging also showed S1B activation by noxious von Frey fibers and laser heat, although the responses to heat trials were smaller compared to those in von Frey trials (likely due to the focused laser beam heating only a small; therefore for fiberphotometry and miniscope imaging studies, we used heating lamp to heat a larger area). S1 may not be the “entry node” for heat signals in cortex, rather S2/IC may be where heat signals first arrive at cortex (*52–54*), the extensive cortical-cortical connections would still allow heat signals to reach S1. We further discovered that while there are many S1B neurons non-specifically respond to all types of sensory stimuli, a subset of S1B L2/3 neurons are only activated in response to a single modality, consistent with the previous evidence in lightly anesthetized primates and rodents (*55–57*). These modality specific S1B neurons, together with other S1B activity patterns (such as the ratio of active neurons, and duration of stimulus-related signals), enables S1B to discriminate the types of sensory stimuli. The signal separation algorithm can be applied to other behaviors with mixed sensory and motor components.

From our single neuron analyses, we found that whisking had a moderate effect on the overall activeness of S1B L2/3 neurons, i.e. whisking further decreases the percentages of active trials of individual neurons in response to noxious stimuli. This finding is consistent with previous studies showing that whisking inputs onto L4 of S1B mediates feedforward inhibition of L2/3 neurons(*18, 24*), although previous studies focused solely on tactile stimuli. This inhibition could partially contribute to the whisking mediated analgesia. On the other hand, whisking had no obvious effect on the highest signal intensities of active neurons in response to noxious stimuli (it only slightly increased the lowest signal intensities). In fact, we observed strong S1B neuron responses to innocuous air puff to whisker and during wiping (as expected). These results suggest that the maximum high intensity of S1B neurons is unlikely a good indicator of pain versus no pain sensation. This idea is completely in line with human EEG and fMRI studies. Human studies showed that the correlation between the magnitude of brain activity in any of the pain-processing cortical areas (S1, S2, IC, and anterior cingulate cortex) and the subjective pain is poor, and can even be dissociated in opposite directions, i.e. there could be prominent activity when no pain is experienced, or there could be small amplitude activity when intense pain is experienced (*58, 59*).

Thus, we examined the population level neural dynamics of S1B in order to extract patterns that are correlated with pain versus no pain responses, and the role of whisking. Using TCA, which is essentially fitting a gain-modulated linear network to the observed S1B activity, we found that larger TCs prior to the expression of nocifensive responses are consistently associated with the subsequent wiping behaviors. While we use wiping as a surrogate for pain being perceived, our finding is consistent with previous studies uncovering choice/decision-related signals in S1B using other behavioral paradigms (*21, 48*). In this regard, S1B can be considered as a region for sensorimotor transformation. The topographic organization of S1B can provide precise spatial information needed to generate a motor plan for an effector (a limb) to target the location (whisker pad). In this framework, the role of whisking could be altering the sensorimotor transformation process thereby leading to a different action (i.e. no wiping in our case). Indeed, by projecting the high dimensional S1B neural activity in each session onto a lower dimensional intrinsic manifold (using manifold learning methods with metric recovery), we were able to calculate the trajectories of S1B population activity in different trials. Despite the fact that the intrinsic manifolds we found from the S1B imaging data might vary across days and across animals, likely due to the variations in experimental conditions (e.g. changes in focus plane, differences in GCaMP expression levels, and low resolutions of miniscope imaging), we nevertheless found that whisking consistently moved the trajectory further away from wiping while shortens the trajectory to the no-wiping action outcome in all cases. Future experiments with larger dataset and higher spatial and temporal resolution imaging may reveal a more salient manifold for S1B neural activity.

In summary, our mouse behavior paradigm provides a new awake animal model to study the neural mechanisms underlying vibrotactile analgesia and more generally, the interactions between touch and nociception. Such studies can now be expanded to other cortical and subcortical regions that we have not investigated in this study. Furthermore, it has been increasingly recognized that the repetitive behaviors observed in autistic patients (often referred to as “stimming”) have a soothing effect for these patients, but the neural mechanisms remain vague. Stimming generates vibrotactile reafferent inputs. With numerous mouse models of autism available, it will be interesting to examine how whisking alters nociceptive processing in both subcortical and cortical levels in these autism models, and perhaps aid future development of devices that can mimic the calming effect of repetitive stimming.

## Methods and Methods

### Animal statement

All experiments were conducted according to protocols approved by the Duke University and MIT Institutional Animal Care and Use Committee.

### Animals

Adult (8-12 weeks) C57BL/6J, Ntng1-Cre (*34*), or Ai162; slc17a7-Cre mice (JAX # 031562; JAX # 023527) were used for the experiments. Animals were housed in the vivarium with a 12-h light/dark cycle and were given food and water ad libitum. Mice were singly housed after GRIN lens implantation.

### Viruses and reagents

AAV2/8-CAG-Flex-GFP (Addgene #59331), AAV2/1-CAG-Flex-GCaMP6m (Addgene #100839), and AAV2/8-Flex-PSAM^L141F^-Y115F-GlyR-IRES-GFP (Addgene #119741) viruses were used for VPM studies. AAV2/1-Synapsin-GCaMP6f (Addgene #100837), or AAV PHP.eB.syn.jGCaMP7f (Addgene # 104488) were used for S1 barrel cortex imaging studies. PSEM 308 hydrochloride was purchased from Tocris (Cat.No.6425) (*60*), and administered (i.p., 5 mg/kg) at 20 min before the behavior test.

### Stereotactic surgical procedures

Mice were anesthetized with isoflurane (3% induction, 1% maintenance) and placed on a digital stereotaxic frame (David Kopf Instruments) with a heating pad. An incision was made in the scalp, and tissue was gently removed to expose the skull. The skull surface was treated by dentin activator (Parkell) before proceeding the following procedures.

#### For viruses expressed in the VPM

small craniotomies were made on the brain surface at the appropriate coordinates (right VPM: +1.95 mm AP, +1.85 mm ML, −3.20 mm DV), viruses (0.3-0.6 μl) were infused at a rate of 50 nl/min using a glass micropipette connected to a Hamilton syringe by tubing filled with mineral oil and left for 10 min after infusion to allow for diffusion.

#### For fiber photometry recording in the VPM

fiber optic cannula (200 μm core, 0.39 NA, RWD) was implanted 200 μm above the VPM after virus injection and secured to the skull using super glue (LOCTITE).

#### For behavior tested on the treadmill

horseshoe shaped headposts were placed on the skull, and Metabond (Parkell) was applied to secure the headpost to the skull, dental cement darkened with carbon powder were used to cover the skull over the top surface.

#### For intrinsic imaging in the right S1 barrel cortex

a small ink marker for the target (right S1 barrel cortex, +1.5 mm AP, +3.5 mm ML) was made on the skull. Headpost was then secured to the skull, and dental cement darkened with carbon powder was used to cover the skull except for the 4-9 mm^2^ window above the barrel cortex. After the dental cement dried, the barrel cortex window was cleaned and filled with saline, which can visualize the surface blood vessels. UV curable adhesive was applied quickly after removing saline with a cotton swab to visualize blood vessels for intrinsic imaging. Subsequently, mice were kept on the heating pad for recovery and returned to the homecage for further intrinsic imaging.

#### For the GRIN lens implantation in the S1 barrel cortex

after aligned location of C2 barrel with cortical vasculature based on the intrinsic imaging data, mice were anesthetized with isoflurane and placed on stereotaxic frame, UV optical adhesive was carefully removed, and a craniotomy (1.5~2 mm ×1.5~2 mm) was made on the brain surface, avoiding damage to the cortex. The craniotomies were shifted medially about ~20° on a horizontal plane for the virus injection. Viruses were (0.6 μl total) infused at a rate of 10 nl/min into 3 locations (−0.25 mm DV) to evenly label the intrinsic signal location. 30 min after virus injection, the GRIN lens (4mm × 1mm, Inscopix) were tightly attached on the virus injected surface and secured with super glue and Metabond (Parkell) darkened with carbon powder. After 3 weeks of virus expression, a holder (Inscopix, gripper part ID: 1050-002199) was used to lower the miniature microscope (Inscopix, nVista 3.0) with the baseplate onto the top of the GRIN lens until the GCaMP6f fluorescence was visible under the illumination from the miniscope’s LED. Subsequently, the baseplate was fixed to the skull with dental cement darkened with carbon powder to prevent external light from contaminating the imaging field of view. A cover (Inscopix, part ID: 1050-002193) was attached to the baseplate to protect the microendoscope.

#### For wide field calcium imaging

mice expressing GCaMP7f (injected retro-orbitally with AAV PHP.eB.syn.jGCaMP7f) were anesthetized with isoflurane (4% induction, 2% maintenance). The scalp and the periosteum were removed and the skull surface and wound margins were covered with Vetbond. Layers of UV-curing optical glue (Norland 81) were applied to reduce light scattering. Metabond was used to secure the wound margins and to attach a metal headpost over the interparietal bone.

### Intrinsic signal optical imaging

Mice were anesthetized with isoflurane (3% induction, 0.7% maintenance) and the whiskers on the left pad, except C2 whisker, were trimmed before starting the intrinsic imaging. A tube for the airpuff connected to a Picospritzer (Parker) was positioned next to the left C2 whisker. Images were acquired with a Basler ace acA1920-155 μm camera with a macroscope composed of 50 mm lens and 35 mm lens. The cortical surface blood vessels were visualized through the intact bone covered with UV-curing optical adhesive. An initial image was taken of the cortical vasculature under green light (525 nm) to enable alignment of intrinsic signal images with surface vessels. Then the camera was focused down below the surface (~500μm). The green light was switched to red 630 nm light for functional imaging, then the images (480 pixels × 300 pixels, 8 mm × 5 mm) were acquired at 10 fps using the custom-written LabView software (National Instruments, TX). Airpuff stimuli delivered to the C2 whisker were applied at 6 Hz for 4 seconds and the intrinsic signal was quantified as the difference in the reflected light during stimulus compared to that immediately before. C2 barrel responses were aligned to the blood vessel pattern to guide the surgery for the craniotomy and viral injection.

### Wide field calcium imaging

Mice were head-fixed on a treadmill and their dorsal cortex was imaged during laser (n = 6; wavelength: 1450nm, power: 300mW, beam diameter: ~1mm, pulse duration: ~200-300ms, 15s interstimulus interval and 20 repetitions) and von Frey stimulation (1g; n = 5; 1s duration, 12-16s inter stimulus interval, 40 repetitions) of the whisker pad. Images were acquired by a PCO.edge 4.2 camera (40 fps, 15 ms all lines exposure time, 500 × 500 pixels, 13 ×13 mm) mounted on a tandem-lens epifluorescence macroscope (front lens: Canon 85 mm F1.8, back lens: Nikkor 105 mm F2). The cortex was illuminated by alternating 405 nm and 475 nm illumination at 20 Hz. GCaMP fluorescence was filtered by a 495 nm dichroic mirror (Chroma T495lpxr-UF2) and a bandpass emission filter (Chroma ET525/50m). Rescaled dF/F_0_ responses measured at 405 nm were subtracted from the dF/F_0_ responses measured at 475 nm to correct for hemodynamic artefacts(*61, 62*). We simultaneously recorded behavior by illuminating the animal with IR light and capturing frames at 200 fps using a Basler acA720-520 um mounted with a UV/VIS cut-off filter (Edmund optics). Synchronization of LEDs, video frame exposure and stimulus timing was controlled by an Arduino microcontroller, using the PCO camera TTLs as a master clock. Behavior video frames were recorded with Bonsai (bonsai-rx.org). Wipe trials were identified using DeepLabCut (*63*). The position of the barrel cortex was determined in separate experiments by mapping the responses to the stimulation of individual whiskers.

### Histology

Animals were deeply anesthetized with isoflurane and then transcardially perfused with pre-cold PBS, followed by 4% pre-cold paraformaldehyde (PFA) fixation solution.

#### For checking the virus expression in the VPM

Whole brains were removed, post-fixed in the 4% PFA overnight at 4 °C, then cryoprotected in 30% sucrose for 2-3 days, frozen in O.C.T compound (Tissue-Tek, Sakura). Then 80 μm coronal brain slices were sectioned using a cryostat. Brain slices were rinsed with PBS between steps and stained with DAPI (Sigma-Aldrich, D9542) for 1 hour, then mounted on the slides for taking confocal images.

#### For checking the virus expression in the S1 barrel

Whole brains were removed, and the cerebral cortex was carefully dissected away from the rest of the brain, flattened and fixed between two slides with 2 mm double-sided tape. And the whole slides were post-fixed in the 4% PFA overnight at 4 °C, then cryoprotected in 30% sucrose for 2-3 days, frozen in O.C.T compound (Tissue-Tek, Sakura). Then 80 μm horizontal sections were cut from the flattened brains using the cryostat. Sections were rinsed with PBS between steps and incubated with primary antibody anti-VGluT2 (Millipore, AB2251-I) overnight at 4 °C, then incubated with secondary antibody Alexa Fluor 647 AffiniPure Donkey Anti-Guinea Pig (Jackson Immunoresearch laboratories, 706-605-148)/DAPI overnight at 4 °C, washed with PBS and mounted on the slides for taking the confocal images. The original confocal images from the same brain cortex included different barrel units, so they were reconstructed and overlapped in photoshop based on their conjunction and structure. Finally, the stacked image showed the virus expression on the complete barrel cortex.

### Behavior experiments

We adapted the mice with headpost were adapted on the running wheel for 2-3 days. Mice underwent behavioral testing sessions with full whiskers and tested again in no-whiskers (whiskers were trimmed) sessions. For the experimental sessions, we applied innocuous (airpuff) and/or noxious (radiant heat and von Frey filaments) stimuli to the mouse whisker pad. For the airpuff experiment, we manually applied the air using a blaster, which typically lasted up to 1 second. For the heat experiment, we used the digital controlled halogen lamp with various percentages of the max intensities and either 4- or 5-second stimuli delivery period (Plantar Test, Hargreaves method, IITC Life Science). We tuned the experimental intensity range covering from no nocifensive to full nocifensive behavior from the mice. For the von Frey experiment (Semmes-Weinstein Von Frey, Stoelting), we also used various grades of filaments (0.02g to 0.4g). The inter-trial intervals for all the experiments were all between 20s to 30s.

#### Initial behavior experiment in Fig. 1

We only tested the noxious stimuli with full whiskers, contralateral whiskers, and no-whiskers. The heat intensities we used were 10%, 15% and 20% of the max intensity of the halogen lamp, and the von Frey filaments were 0.02g, 0.04g, 0.07g, 0.16g and 0.4g. For each intensity, we repeated the trials 10 to 20 times on n = 9 (5 male and 4 female mice, for both full and no whiskers experiments) C57BL/6J mice, and n = 7 (3 male and 4 female mice, for contralateral whiskers experiment).

#### Fiber photometry experiment in Fig. 3

Fiber photometry recording (RWD) was used to record the population calcium fluorescence from the VPM_Ntng1_ neurons. We applied both innocuous and noxious stimuli with full whiskers. We chose the optimal intensity for the noxious stimuli (heat 15%, von Frey 0.4g) that elicited nocifensive behaviors and also allowed whisking induced analgesia. Individual stimuli were repeatedly applied 10 to 20 trials on the whisker pad of VPM_Ntng1-GCaMP6m_ mice (n = 6), video and calcium signal were simultaneously recorded.

#### Manipulation experiment in Fig. 4

We applied only noxious stimuli with full whiskers. The heat intensities we used were 10%, 15% and 20%, and the von Frey filaments were 0.07g, 0.16g and 0.4g. We repeated the trials 10 to 20 times on the whisker pad of VPM_Ntng1-PSAM_ mice (n = 8). For each stimulus type per mouse, we conducted a pre-drug and a post-drug session as baseline controls, in addition to the PSEM308 session which blocks the VPM_Ntng1_ input signals to S1.

#### Calcium imaging experiment in Fig. 5 and 6

We used Inscopix nVista3.0 system to record the calcium dynamics from the neuronal population of S1B L2/3. The frame rate of all the sessions was 20 fps. We applied both innocuous and noxious stimuli with both full whiskers and no whiskers. Again, we chose the optimal intensity for the noxious stimuli (heat 15%, von Frey 0.4g) that elicited nocifensive behaviors and also allowed the whisking induced analgesia. For each intensity, we repeated the trials ~30 times (innocuous), and 60 to 120 times (noxious) on C57 mice (n = 6), and recorded the calcium signals of L2/3 neurons of S1B C2 and neighboring barrels.

### Behavioral data analysis

All behavior videos were manually processed. We tracked stimulus onset moments, whisking periods and wiping moments throughout the whole video. Specifically, stimulus onset and whisking on/off moments were tracked precisely to a single frame, and for the wiping, each time the paw touching the whiskers/face was considered as a wiping moment. For the no-whisker trials, we labeled the whisking periods based on mice facial muscle movement. Additionally, throughout the experiments, we found the coupling between the mouse whisking and running was high (data not shown). The tracked factors were then converted into a binary labelling indicating the framewise on and off of each factor. Moreover, to include the potential lasting effect of the stimulus and wiping, a duration of effect was assigned to each stimulus onset and wiping moments. For airpuff and von Frey (often with a single wiping) trials, the duration of effect was 2 seconds, while for heat trials, the duration was 5 seconds.

For the behavioral analysis (initial behavior in Fig. 1 and VPM_Ntng1_ chemical slicing experiment in Fig. 3), two measures, wiping numbers (how many times a mouse wiped the face during a trial) and wiping trial percentage, were used to characterize the level of the nocifensive behavior. The mean and standard error were calculated by the wiping number of all the trials, while the wiping trial percentage was calculated by that of the individual mice.

### Fiber photometry data analysis

#### Signal preprocessing

The 470 nm signal was baseline corrected by the 410 nm channel, and photobleaching was corrected by adaptive baseline estimation custom written in MATLAB. For the baseline estimation, the signal was first cleaned using wavelet signal denoising. Then the key turning points were extracted by Ramer-Douglas-Peucker algorithm (*64, 65*), followed by adaptive thresholding (see below) to find the key turning points representing the baseline dynamics. The baseline was then estimated by piecewise linear interpolation using the key baseline turning points.

#### Factor related signal extraction

We assume that the factor related signals follow the rule of linear superposition. The signal was first converted to deconvolved calcium signal space using constrained foopsi (*36*). The deconvolved calcium signal could be considered as rescaled overall calcium influx from the neural population that was being recorded. Then all the labelling we extracted from the behavior videos for the three factors, stimulus (either airpuff, or heat, or von Frey), whisking and wiping, were combined to be a 3-fold factor labelling (the factors of stimulus, including airpuff and von Frey, and wiping were attached with a window of ~1 second to include the potential lasting activity), e.g. [1, 1, 0] indicating this frame contains stimulus, whisking but not wiping. Each frame belonged to one of the 2^3^ = 8 conditions. To extract one factor related signal, we first collected all the calcium imaging frames with this factor labelling being 1, and then pooled frames based on the type of the labelling, for example, [1, 1, 0], [1, 0, 0], [1, 0, 1] and [1, 1, 1] are types of conditions containing the stimulus factor. For each condition, we calculated the intensity distribution of the deconvolved signal, and then calculated the distribution of the corresponding factor-negative conditions, for example, [0, 1, 0] and [0, 0, 0], [0, 0, 1] and [0, 1, 1] for the above example. By assumption, the distribution difference between conditions only differing in the presence or absence of the factor of interest could be approximately ascribed to the pure factor related signal. After calculating the factor related distribution, we then applied a randomized event pick procedure to generate the pure factor related deconvolved signal. For each intensity subrange of the distribution, we randomly selected the number of frames based on the factor related distribution whose deconvolved signals were falling within the subrange. We repeated this process multiple times (n = 100) and then computed the average of the realizations. This average was then considered as the pure factor related deconvolved signal. We applied the method to whisking and wiping separately, and generated the stimulus related factor by calculating the residual of subtracting the above two signals from the raw signal to compensate for the fewer time points for the stimulus factor. To calculate the fluorescence signal, the deconvolved signal was then convolved using the same parameters of the autoregressive process.

### In vivo calcium imaging data analysis

#### Calcium data processing

The calcium imaging data were processed using MIN1PIPE (*38*). The factor related signal extraction described above for fiber-photometry was extended to multichannel calcium imaging data by picking events from multichannel signals instead of the same single channel, and all three factors were estimated by the method. Post processing manual ROI selection was conducted to remove any non-neuron like ROIs before any analyses, independent of the analysis procedures. For a single behavioral configuration, the combination of key factors of interest such as stimulus, whisking and wiping, we pooled the data from that configuration into a data tensor of the dimensions of neuron number, length of a trial, and trial number. For different average measures, we operated along corresponding dimensions.

#### Calcium data analysis

To extract active neurons or trials, we adaptively calculated the threshold based on each neurons’ n times (n = 1 or 2) of standard deviation. For active neuron ratio, a neuron is considered active during the trial if it had at least 1 calcium event larger than 1 standard deviation of the signal in each trial (trial duration 5 seconds for heat, and 2 seconds for von Frey). For active trial percentage of a given neuron, the threshold was defined by 1 standard deviation of the max intensity of all trials for that neuron. By this criteria, if a neuron has background activity, and if the stimulus did not increase its calcium signals in a trial, then the neuron is not considered having an active trial.

#### Same-cell tracking (across sessions/days analysis)

To track the same neurons across all sessions, we applied modified CellReg (*39*) on all the sessions of the same mouse. Because a direct tracking using one of the sessions as the reference would increase the missing rate when the target session was far apart from the reference, we thus tracked the same neurons in a cyclic repeating pattern. In each repetition, a different session served as the reference session on which other sessions were registered. We first extracted the maximal number of trackable neurons to be the minimum of the maximum neuron number from each repetition. Then for each neuron, the corresponding neuron identity was extracted for each repetition, and the final correspondence was set to be the mode of all the individual corresponding identities.

#### Tensor component analysis

We applied the TCA (*41, 66*) on the fluorescence data tensor to decompose the tensor into individual components. To identify neural correlates of S1B activity with facial nociception, we focused on the collective feature of a group of components, rather than individual components to reduce the potential bias. Specifically, we assigned a large number of expected components to improve the fitting performance (n = 80). We also ran the modeling for each session multiple times (n = 50) and then the decomposed trial factors of all the components were compared with the real behavior factor readouts (i.e whisking, no-whisking, wiping, no-wiping), and a cosine similarity was computed for each of them. The first m = 4 components with the top similarity were extracted from each mouse and formed a matrix of behavior relevant components of size 24. Because in TCA, there is no unique scale information in the decomposition, we thus normalized all the components for the downstream analyses. To calculate the weighted temporal factor for subclusters focusing on pain versus no pain sensed (i.e wiping versus no wiping responses), we first divided the temporal domain into 1-second bins and focused only on the bin containing the neural dynamics right before the wiping behavior ([2, 3s] for heat and [0, 1s] for von Frey). Then we extracted the temporal factors from the matrix of behavior relevant components whose rising phase (for heat; peak for von Frey due to the rapid wiping behavior after stimulus application) fell within the bin range, and computed the average of these factors with the scale of the percentage of the factors in this bin.

#### S1B activity manifolds analysis

A manifold of the S1B activity was learned for each session. First, we formed the data matrix of that session using the deconvolved calcium signal, which was a n × T matrix where n was the number of neurons and T was the number of time points. The data matrix was then binarized through thresholding using 1 standard deviation of each trace, and time points with all zero elements were excluded (*67*). To estimate the intrinsic dimension of the manifold, we applied PCA on the data matrices and considered the first n dimensions whose cumulative variance explained by the n dimensions exceeded 97.5% (Supplemental Fig. 7) (*43*). A manifold learning algorithm, Laplacian Eigenmap, was then applied to the data matrix (*42*), with d = 10 as the dimension of the embeddings, employed based on the initial estimation of the intrinsic dimensions. An adjacency matrix was first constructed by computing the nearest neighbors (p = 0.25% for the first iteration and p = 10% for the second iteration) (*67*). Then a weight matrix was constructed using the parameter free method based on the adjacency matrix (*42*). The problem of manifold learning was then converted to solving an eigenvalue problem. The learned manifold was usually spatially distorted, resulting in the failure of preserving the geometry of the data. Therefore, we applied Megaman (*45*) to recover the geometric information of the learned embedding. The metric learned applied to any pairs of points on the embedded manifold.

To further estimate the embeddings of the subspaces for different behavioral configurations, we applied PCA and preserved the first dimensions which explained 99% variance in total. To extract the shared dimensions between two subspaces, we applied a method using basic linear operations, given the bases vectors of the two input subspaces. To calculate the trajectory length along the embedded manifold, we converted the embedded manifold to an undirected graph, with the distance between pairs of points to be corrected by the learned metric. Then for any neighboring two temporal points along the trajectory, we computed the shortest path (Dijkstra’s algorithm) (*68*) on the graph and calculated the distance of this shortest path, as the approximation of the geodesic length on the manifold. To finally get the trajectory length, we summed up the distances of all the neighboring temporal points along the trajectory within a trial computed in the above way.

### Statistics

All statistical analyses were performed in Matlab (Mathworks). Unless mentioned explicitly, most behavior data, including the full whiskers, contralateral whiskers and no whiskers behavioral experiments were tested with two-tailed paired t-test. Fiber photometry data were tested with two-tailed two sample t-test. Manipulation experiments and wide field imaging data were tested with one-way repeated measures ANOVA, where the former was also tested with post-hoc Tukey’s honestly significant difference examination between pairs of conditions. Further linear mixed-effect model was applied to both heat and von Frey experiment where there was no strong significance due to the information loss at computing the mouse average for ANOVA. For distribution comparison, Kolmogorov-Smirnov (KS) test was first applied to directly test if the two distributions are from the same one. If test returned null hypothesis, it was then followed by bootstrapping measurement of the relative entropy between the distributions, which was further examined by t-test for the mean. Significance levels are indicated as following: *, p < 0.025, **, p < 0.01, ***, p < 1 × 10^-4^ and ****, p < 1 × 10^-6^. Data distribution was assumed to be normal, but this was not formally tested; instead, all graphs contain individual data points and mean ± s.e.m..

## Acknowledgments

We thank Thuy Hua for the initial help with the von Frey experiment, and Jaehong Park for help with the fiber photometry.

## Author contributions

F.W. conceived and supervised the project. F.W., J.L. and B.C. designed the experiments. B.C. and J.T. performed surgeries and virus injections. J.L., B.C. and P.X. performed behavioral tests. M.L. and V.P. built the wide-field imaging system and performed wide field imaging. B.C., M.L. and P.M.T. performed intrinsic imaging. J.L. and B.C. performed calcium miniscope imaging. J.L. processed and analyzed the behavioral and neural data. B.-X.H. took care of mouse husbandry and genotyping. Z.H. provided the critical reagents. F.W., J.L., B.C., M.L. and V.P. wrote the manuscript.

## Data and materials availability

All raw data described in this study are available from the corresponding authors upon reasonable request. All codes described in this study are available from the corresponding authors upon reasonable request. The MIN1PIPE code for processing miniscope imaging (correspondence should be addressed to Jinghao Lu) is available at: https://github.com/JinghaoLu/MIN1PIPE.

