## Supplemental Materials for "Somatosensory Cortical Signature of Facial Nociception and Vibrotactile Touch Induced Analgesia"

**This PDF file includes:**

Figs. S1 to S8
Captions for Movies S1 to S2

**Other Supplementary Materials for this manuscript include the following:**

Movies S1 to S2

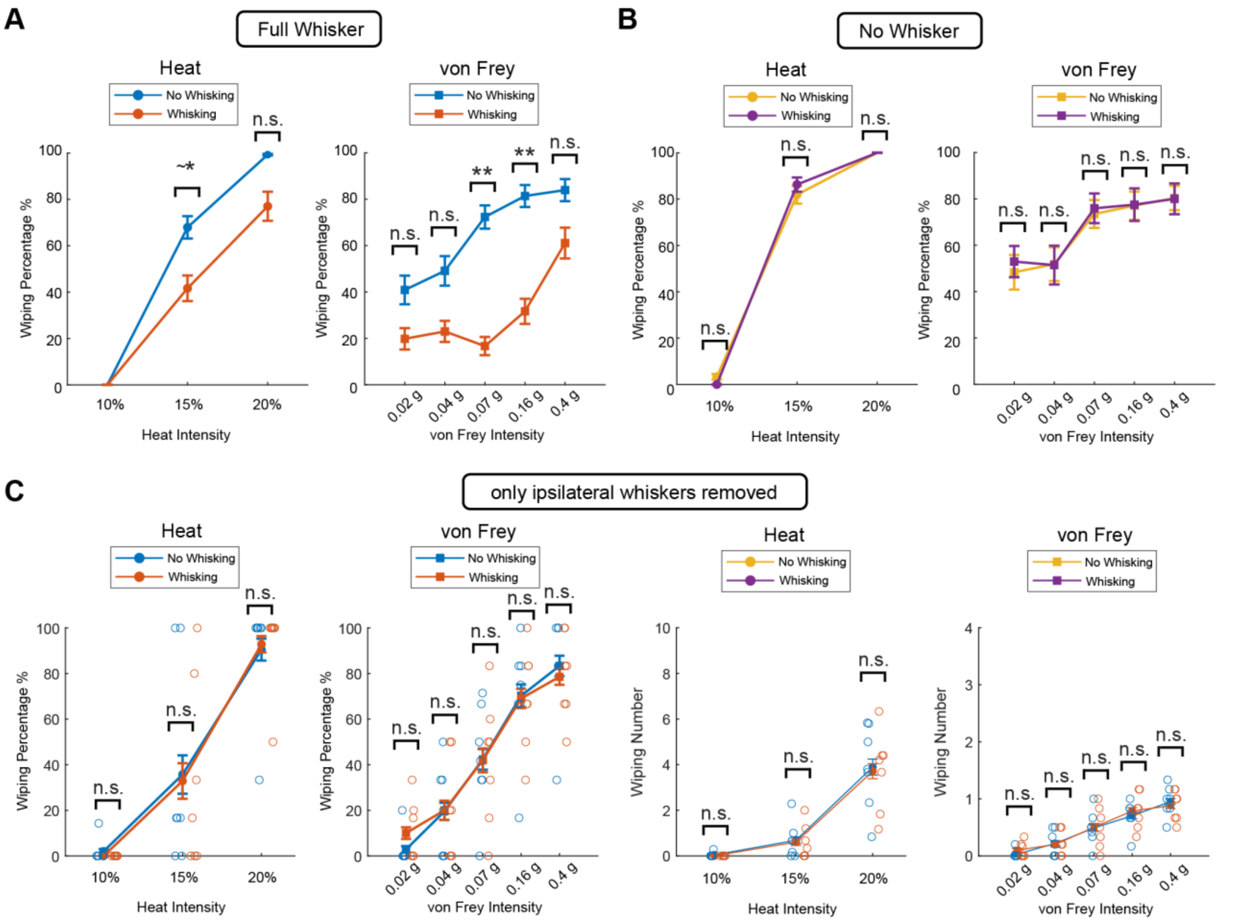


**Fig. S1. Ipsilateral reafferent whisking signal suppresses facial nociception, related to Fig. 1. A-B.** Nocifensive behavior in response to heat or von Frey stimuli as measured by the percentage of trials with wiping behaviors in the full whisker condition (**A**, n = 9) or no-whisker conditions (**B**, n = 9). **C.** Nocifensive behavior in response to heat or von Frey stimuli in mice with ipsilateral whiskers removed while the whiskers on the contralateral stimulation side remain intact (n = 7). Left, measured by wiping percentage. Right, measured by wiping number.

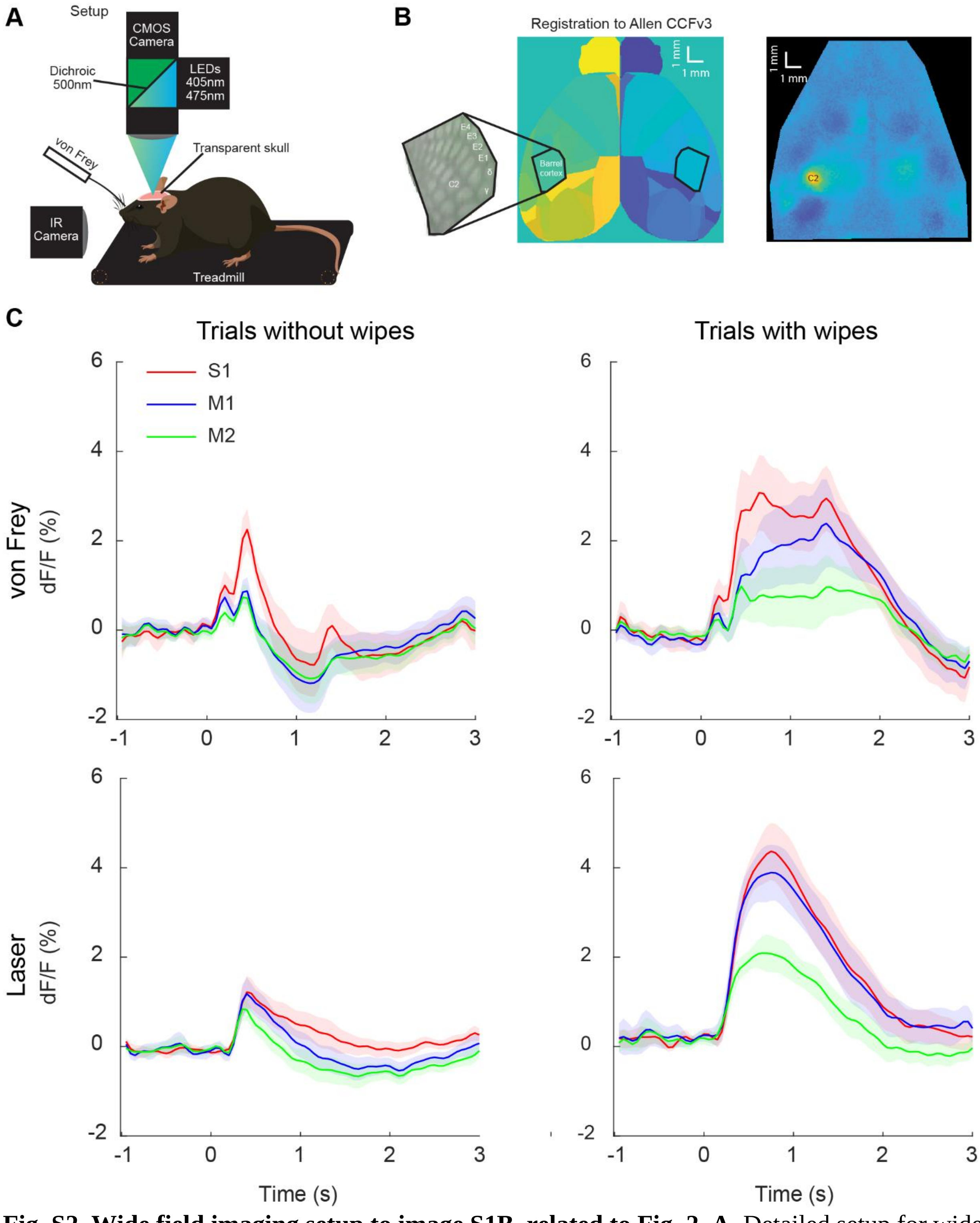


**Fig. S2. Wide field imaging setup to image S1B, related to Fig. 2. A.** Detailed setup for wide field imaging of calcium activity. **B.** Left: Registration to the Allen atlas. Wide field responses to single whisker stimulations are used to localize the corresponding barrels. 2-3 barrels are mapped per hemisphere. Right: Example of response to stimulation of the right C2 whisker. **C.** Averaged calcium fluorescence in S1B, M1 and M2 under different experimental conditions (n = 6 in heat and n = 5 in von Frey). M1, primary motor cortex. M2, Secondary motor cortex.

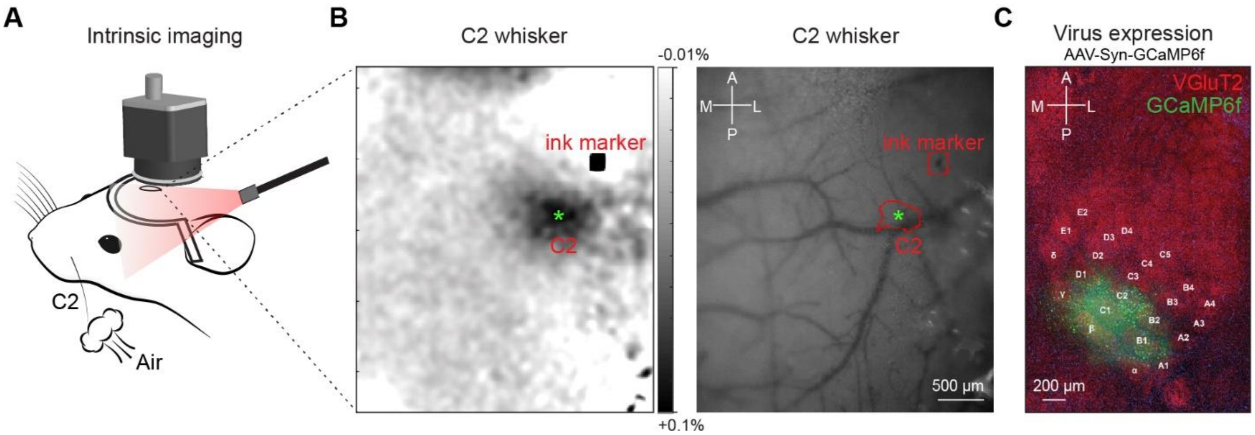


**Fig. S3. Identification of C2 whisker barrel in S1B with intrinsic imaging for targeted injection of AAV-syn-GCaMP6f, related to Fig. 5**. **A.** Intrinsic imaging setup. Airpuff stimulates the C2 whisker of an anesthetized, head-fixed mouse. The camera captures a sequence of S1 cortex images under the illumination of a red (630 nm) light source. **B.** Left, Intrinsic imaging response map, the dark area (green asterisk) indicates the cortical region activated by C2 whisker stimulation. Right, the corresponding blood vessel map obtained with green illumination. The blood vessel map was used for targeted viral injection. **C.** Post hoc histology of GCaMP6f expression in the barrel cortex from the same mouse as in **B**.

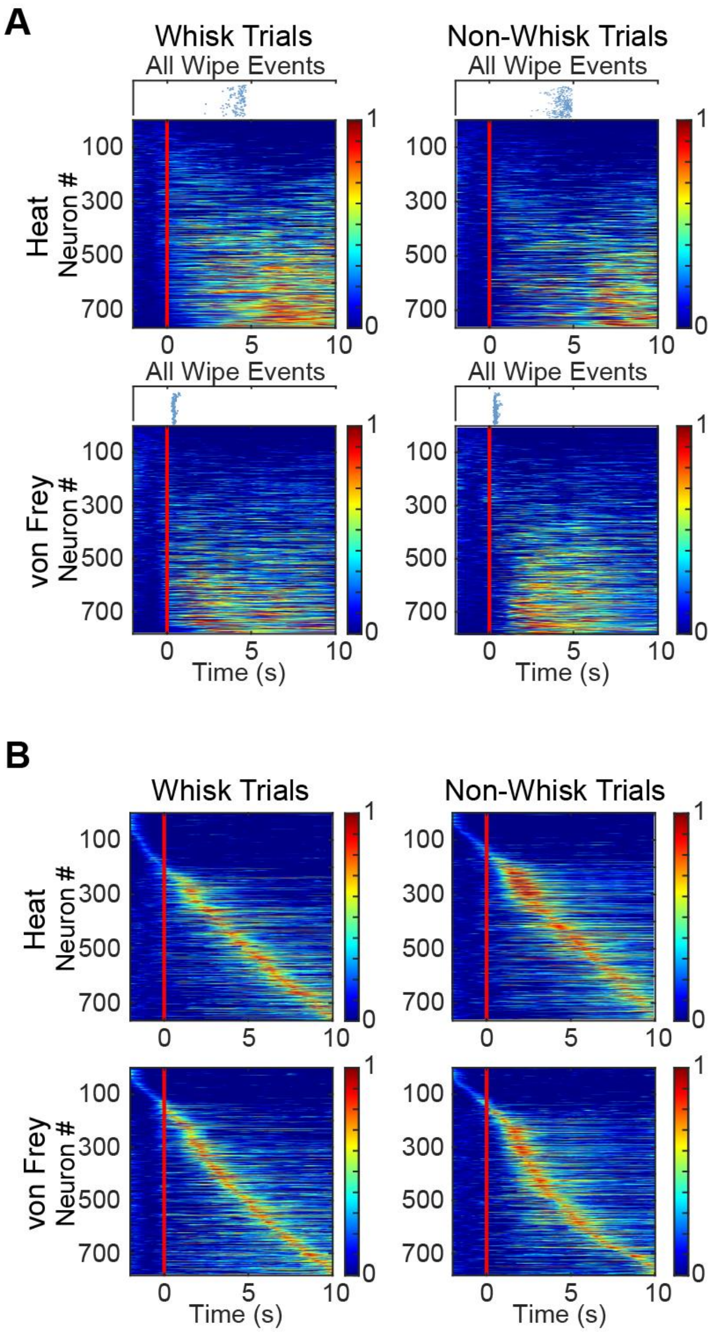


**Fig. S4. Alternative ways of sorting S1B neural signals, related to Fig. 5**. **A.** All trial averaged neuronal traces sorted by the maximal intensity, aligned to the stimulus onset (red line is stimulus onset). Note that the stronger signals appear after wiping events, consistent with wiping whole face generates the strongest tactile stimuli. Mice often wipe multiple times, thus the most intense signals appear with a delay to the initial wiping event. **B.** All trial averaged neuronal traces sorted by the peak latency, aligned to the wiping onset (red line is wiping onset). While signals are stronger after wiping onset (as expected wiping generates the strongest tactile stimuli to face), there are signals preceding the wiping onset, reflecting stimulus (heat or von Frey) related signals.

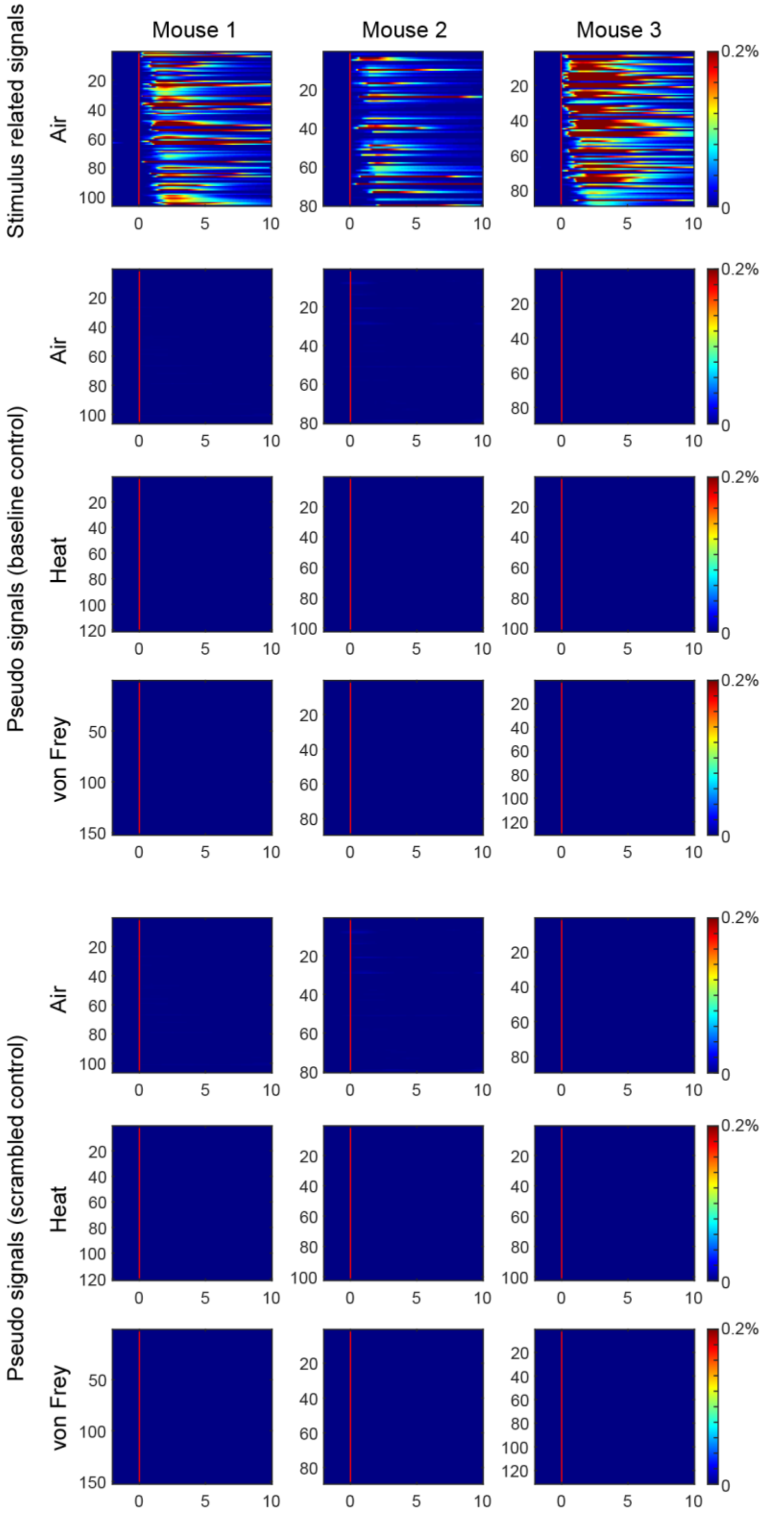


**Fig. S5. Control analysis for pseudo stimulus related signals using the same multivariate generalized factor related signal estimation method, related to Fig. 5.** Trials of pseudo signals of example mice estimated in the experiments of different stimulus types (both baseline control and scrambled trial control; see Methods for details). Top row consists of stimulus related signals in the airpuff stimulus experiment, shown here as a reference.

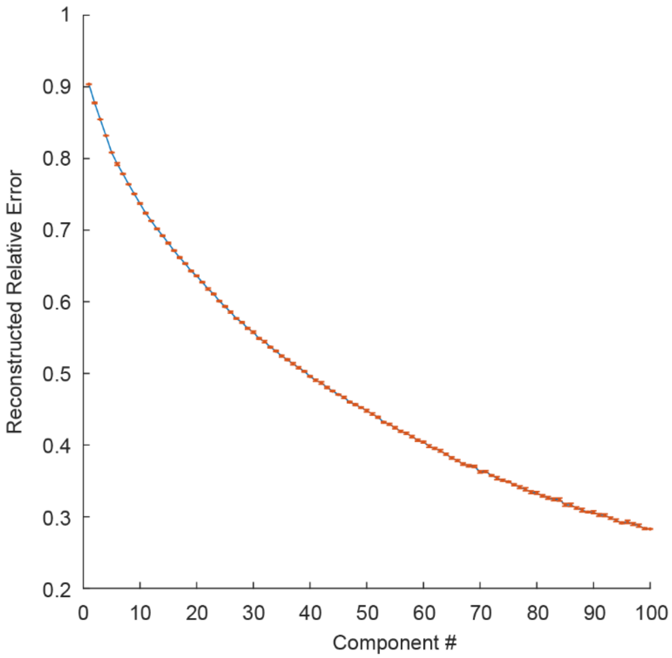


**Fig. S6. Reconstructed relative error of TCA modeling as a function of numbers of components used as inputs, related to Fig. 6.**

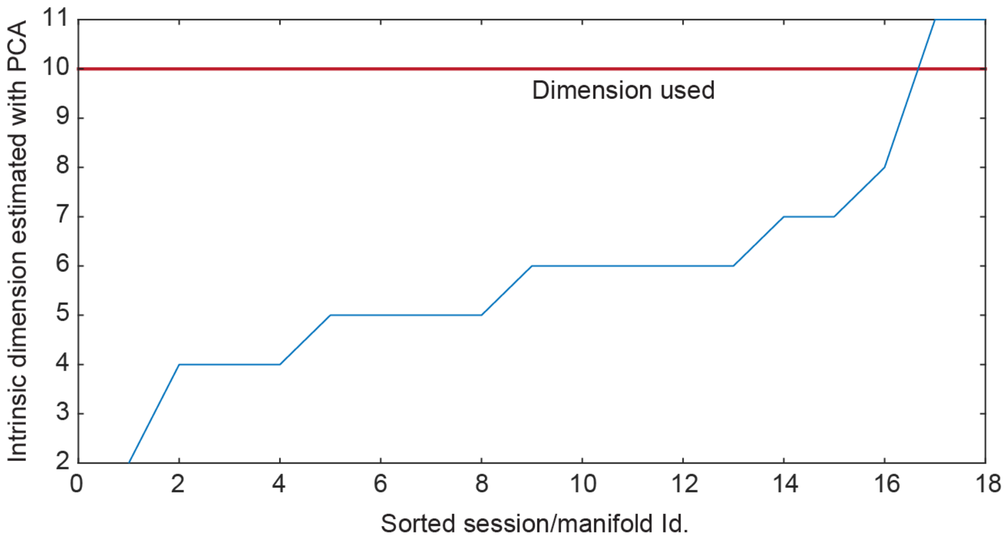


**Fig. S7. Numbers of intrinsic dimensions of S1B neural activity from individual sessions estimated using PCA, related to Fig. 6**. The estimated dimensions were sorted in increasing order, and based on the most sessions, a global intrinsic dimension of 10 was used in the later manifold learning.

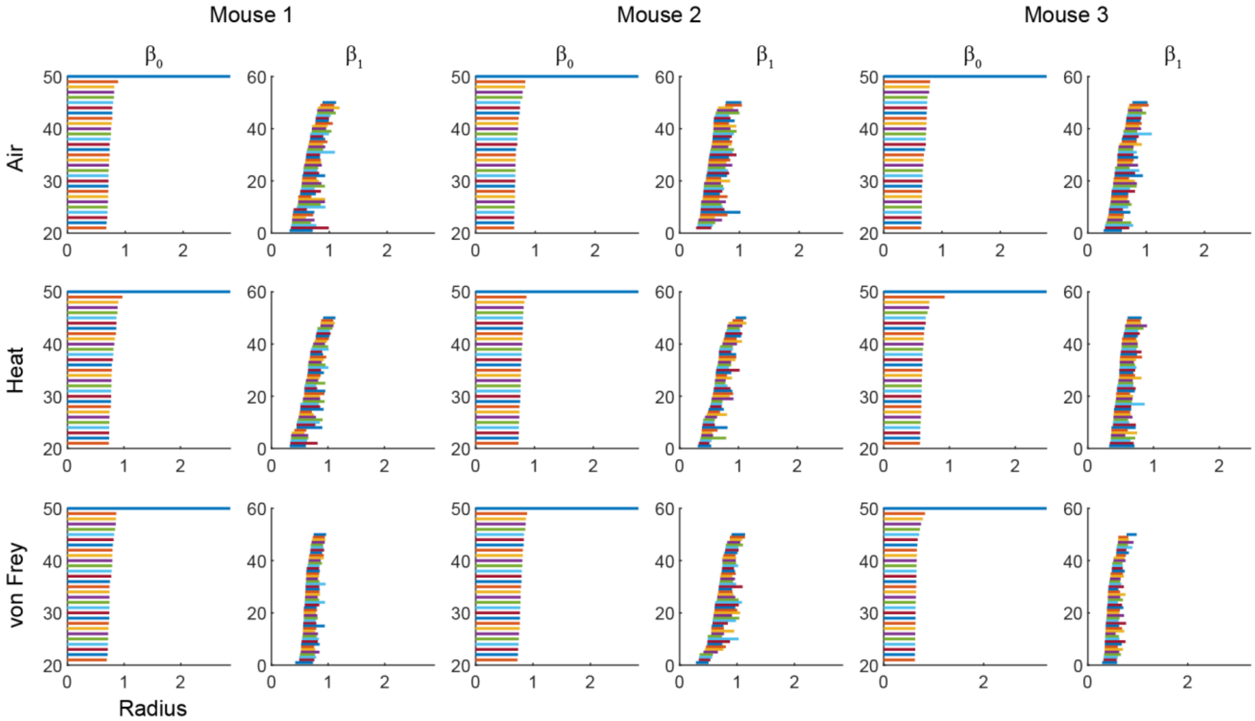


**Fig. S8. Persistent diagrams of the learned intrinsic manifolds in three example mice, related to Fig. 6.** Diagrams showing the Betti numbers up to 1 dimension are included, due to the time consuming computation.

**Movie S1.**

**Behaviors showing whisking induced analgesia behaviors.** Examples of the four types of behaviors in response to mechanical von Frey stimulus: full-whisker + whisking + no-wiping (top left), full-whisker + no-whisking + wiping (top right), no-whisker + whisking + wiping (bottom left), and no-whisker + no-whisking + wiping (bottom right).

**Movie S2.**
**S1 barrel cortex responses to von Frey stimuli.** Example trials of wide field calcium imaging signals of the S1 barrel cortex in response to the applied von Frey stimuli.
